# *parSEQ:* Probe and Rescue Sequencing for Advanced Variant Retrieval from DNA Pool

**DOI:** 10.1101/2023.12.12.571337

**Authors:** Moustafa Houmani, Finlay Peterkin, Gerard Antoun, Louis Fischer, Anissa Hammi

## Abstract

Modern protein engineering is powered by sequence-function data-sets. We have developed parSEQ, a platform that maximizes the capture of these protein sequence-function data-sets through a sequence-first-screen-later approach. parSEQ relies on Next-Generation Sequencing (NGS) to reverse the conventional screen-first-sequence-later workflow. This allows for the high-throughput retrieval of variant sequences from DNA pools, ensuring that every screened variant’s functional data is paired with its sequence data. This report details parSEQ’s methodology and its integration into various protein engineering workflows. Through several case studies, we illustrate parSEQ’s broad applicability. These case studies describe the use of parSEQ for high-throughput variant DNA-template retrieval, expression-ready bacterial clone isolation, sourcing DNA for denovo designed proteins, and preparing targeted mutational libraries. Our findings suggest that parSEQ’s approach to capturing sequence-function data-sets can advance protein engineering efforts, particularly in the age of artificial intelligence and machine learning.

## 1 Introduction

Proteins are fundamental building blocks of life, essential to the majority of biological functions. Humans have long recognized the significance of proteins and have greatly expanded their use beyond their natural functions. Today, the exploitation of proteins is broad in scope, from therapeutics to industrial catalysts and diverse biomaterials. ^1,2,3^. Despite this, the challenge of engineering proteins remains a complex experimental process, managed by a select few organizations globally. The creation and development of new proteins can be conceptualized as a search problem, a search for the optimal sequence of amino acids that will fold into a functional protein with the desired characteristics and biological activity. Today, researchers use Machine Learning (ML) and Artificial Intelligence (AI) tools to guide this search problem because these tools can navigate the sequence-function landscape of proteins more efficiently ^4^. AI and ML tools can sift through vast amounts of data, analyze patterns, and suggest the most promising amino acid sequences to satisfy the desired characteristics and biological activity. The effectiveness of such tools, however, is intrinsically linked to the quality and volume of experimental data-sets ^4,5,6^.

Sequence-function data-sets are a cornerstone for engineering proteins, where structure is also considered a facet of function ^7,8^. A thorough understanding of how sequence alterations impact protein function can illuminate foundational biological mechanics and inform predictive modelling and protein design. ^9^ Given this, maximizing the capture of sequence-function data-sets within time and budgetary constraints is essential for any protein engineering effort ^7,10^.

Unfortunately, the capture of sequence-function data-sets is impeded by the prevailing use of the screen-hit-sequence workflow in many contemporary protein engineering workflows ^11^. For instance, an advanced protein binder development program typically involves high throughput binder filtering (display methods), colony picking, protein binder expression, binder affinity screening (using ELISA, BLI, SPR…), followed by the sequencing of top contenders ^12^.

This blind approach, though entrenched, is primarily driven by the economic constraints associated with Sanger sequencing, the gold standard of variant sequencing today. Sanger sequencing works perfectly well for such an application, but its cost scales linearly with the number of variants sequenced. This cost structure is the reason behind the screen-first-sequence-later approach, as it aims to reduce the number of variants sequenced using screening as a filter. However, such approach ultimately results in limited hits and sparse data (Figure 1). Moreover, post-hit sequencing analysis often reveals that the sequenced variants are not as diverse as anticipated (multiple identical hits), restricting downstream experimental flexibility.

**Figure 1:**
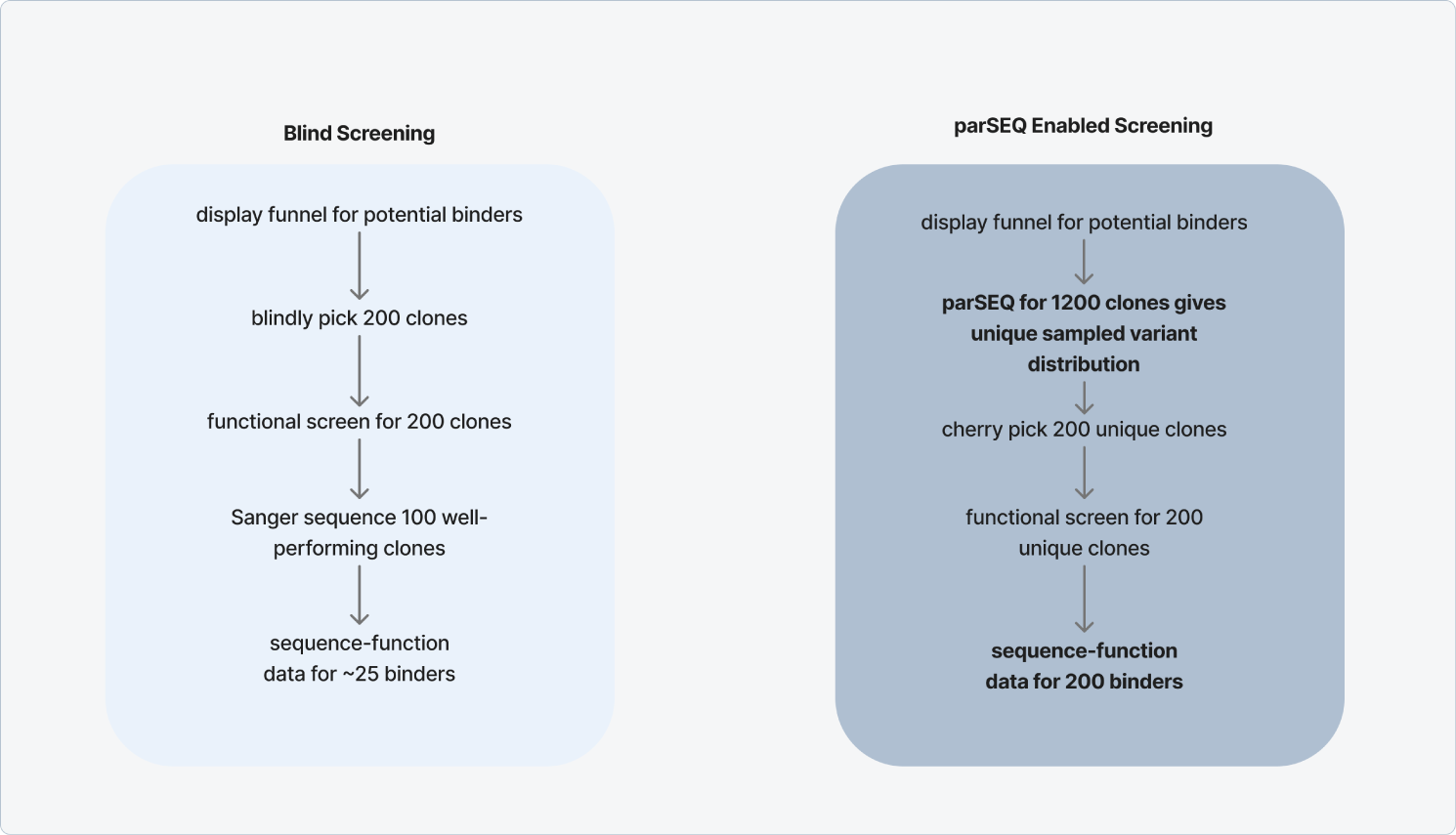
Blind screening vs parSEQ-enabled screening approach. parSEQ’s sequence-first-screen-later approach yields considerably more sequence-function data for the same cost. In traditional blind screening approaches (screen-first-sequence-later), the number of clones picked is limited by the screening capacity and downstream Sanger sequencing cost. In the case of a display pool output of protein binders, and with a binding screening capacity of 200 binders, 200 clones of the display pool output will be blindly picked. This is followed by binding screening (e.g. using BLI) that yields 100 well-performing clones. Sanger sequencing on these 100 well-performing clones reveals only 20-30 unique binders due to the pool enrichment profile, compounded by the additional enrichment of screening for well-performing variants. This ultimately results in sequence-function data for 25 unique variants. parSEQ allows researchers to choose their variants before going for functional screening using a sequence-first-screen-later approach. Under such paradigm, parSEQ ingests the display pool output and reports an enrichment profile of the sampled unique variants in the pool, after having physically separated these variants into clonal wells. A researcher can then cherry pick 200 unique variants to go for functional screening. This ultimately results in sequence-function data for the 200 cherry-picked unique variants. This is 8 times more sequence-function data for the same cost and under the same timelines (costs and timelines will be discussed in more detail below).

We perceived a significant advantage in reversing the conventional screening-sequencing order. Adopting a sequence-first-screen-later methodology facilitates a more expansive understanding of the protein sequence-function landscape (Figure 1), because it ensures that we collect sequence data for every screened protein variant’s functional data. Moreover, it ensures that we never screen the same variant more than once.

To that end, we developed parSEQ, a platform engineered to handle various DNA variant pools, delivering sequence-verified, physically separated variants. The process (Figure 2) begins with the distribution of bacteria from collective pools into separate clonal wells. After this, parSEQ employs a two-level barcoding system for enhanced multiplexing, incorporating both well-specific and plate-specific barcodes. Once barcoded, samples are pooled for Next-Generation Sequencing (NGS), after which the NGS data is analyzed to link each variant’s sequence with its respective well. Operating within a 384-well plate setup, and supported by automated procedures and Python-based data analysis, parSEQ can sample variant pools with high efficiency and fast turnaround times. While initially dependent on Illumina sequencing for accurate sequencing results, Oxford Nanopore (ONT)’s recently developed v14 chemistry & R10.4.1 pore, along with improved basecalling algorithms now position ONT sequencing as an effective substitute ^13^.

**Figure 2:**
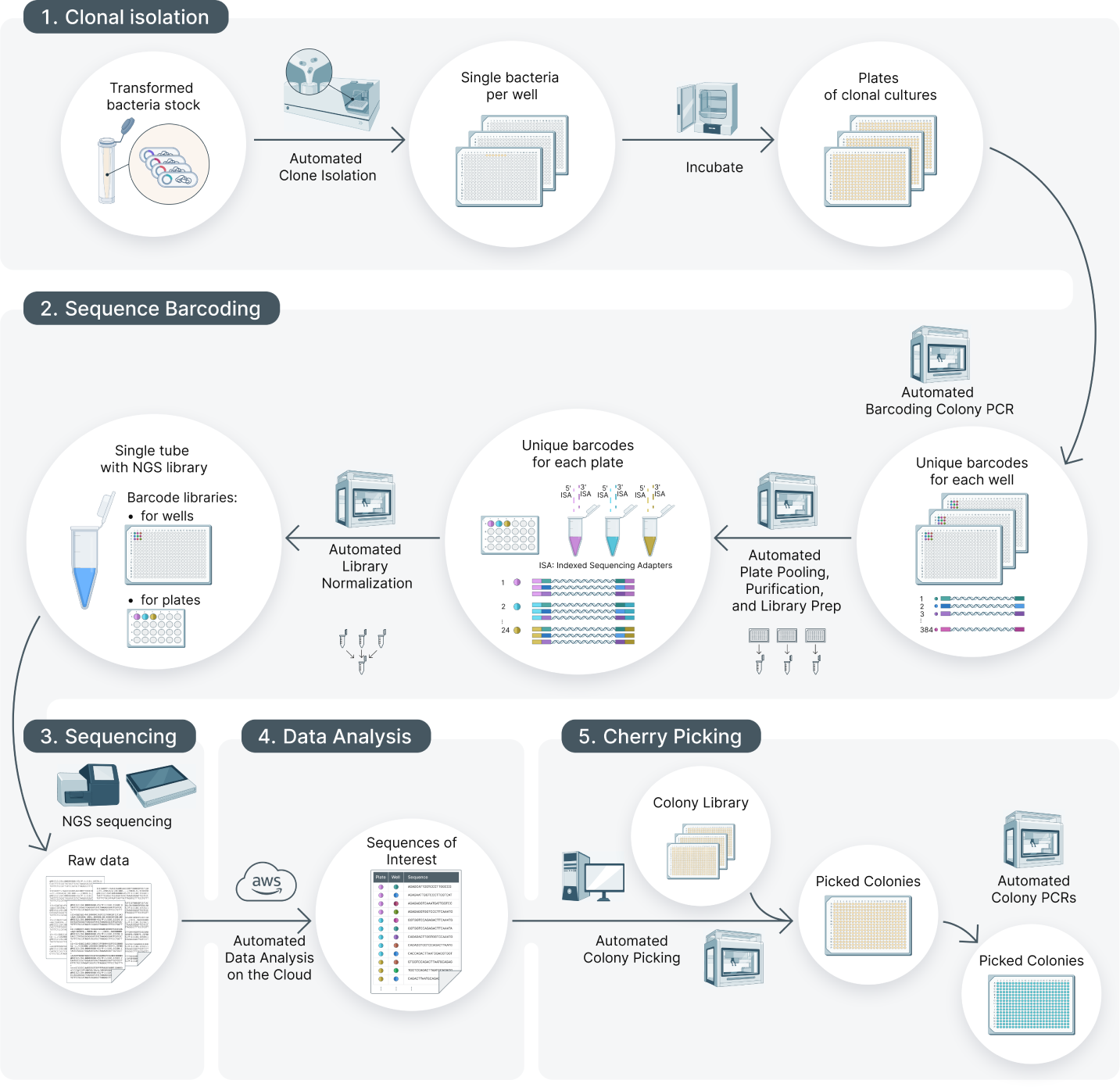
parSEQ process workflow **Clonal Isolation:** Utilizing a single bacterial cell dispenser like Cytena’s B.Sight, individual bacterial cells from a bacterial pool are allocated into wells of 384-well plates. Following a 24-hour incubation period to permit bacterial growth, each well contains only one bacterial clone. It is crucial to clarify that each well may not necessarily contain a unique DNA sequence (i.e. the same sequence will probably exist in multiple wells), as the sampled wells collectively reflect the diversity and enrichment within the original pool. Clonal isolation can thus be thought of as **sampling from the original pool**. **Sequence Barcoding:** Employing a liquid handler such as the Hamilton Starlet for rapid high-throughput PCR reaction prep, we run barcoded colony PCRs to barcode and amplify the DNA sequences within each well of each plate. Subsequently, the PCR products from each plate are pooled into a tube and purified using AMPure magnetic bead-based purification, yielding a pool of barcoded PCR products for each plate (each plate goes into one tube, no plate-level mixing yet). Each barcoded PCR product pool undergoes an NGS library prep, incorporating barcoded adapters distinctive to each pool (and consequently, each plate). This ligation-based library prep avoids the necessity for any amplification steps, eliminating the risk of variant recombination. Post library-prep, the samples of the different plates are pooled into one sequencing ready pool. **Sequencing:** The consolidated libraries are sequenced on the chosen NGS platform. **Data Analysis:** The obtained NGS reads are demultiplexed at the plate level using the NGS adapter barcodes. Subsequently, the reads within a plate are allocated to their respective wells via the PCR barcodes (well-based demultiplexing). Well based sequence alignment is performed, and the aligned reads for each well are then analyzed to derive each well’s consensus sequence. After retrieving the well-based consensus sequences, the unique sequences are tallied, and the sequences of interest are selected. **Cherry Picking:** The wells with sequences pinpointed as being of interest are cherry-picked from the original plates and cultured in new 384 well plates. At this point, users have acquired sequence-verified clones in the wells of 384-well plates, ready for any further applications or analyses they intend to pursue.

With parSEQ, users have the capability to:

*⋄* Barcode up to 96×384 selected clones (96x 384-well plates), totaling 36,864 clones, in a single run. This limitation arises from plate-level barcoding, dependent on the indexing provided by commercial barcoding NGS adapters (Illumina sequencing adapters, ONT sequencing adapters).

*⋄* Extract variants of interest from the pool. The attainable number of unique variants is contingent upon both the number of wells (or clones) processed and the pool’s enrichment profile.

*⋄* Use the retrieved variants for any characterization assays users wish to conduct.

When we started working on parSEQ, there were no comparable tools reported in literature. In early 2022, evSeq was released by the Arnold lab ^14^. While evSeq shares its foundational concepts and goals with parSEQ, our implementation enhances the efficiency and scale of pooled variant sequencing. parSEQ leverages 384-well plates, automated processes, and cost-effective in-house (ONT) sequencing to achieve higher throughput and rapid turnaround times. We advocate for the integration of parSEQ in the workflows of any team actively engaged in protein engineering, particularly those leveraging machine learning and/or AI. parSEQ empowers users to delve deeper into pools, isolate rare variants, and meticulously map out the sequence-function landscape. In this technical report, we will showcase how parSEQ can be used to retrieve sequence verified & clonally isolated variants from display pools, source DNA templates for denovo designed protein libraries, and help retrieve sequence verified & clonally isolated variants for targeted mutational libraries.

## 2 Methods

In the methods section, we first provide a detailed description of the parSEQ process workflow. Next, we explain how labs of different sizes can adopt parSEQ, even with limited financial resources, emphasizing the low capital expenditure required. We then present a practical breakdown of the costs and timelines. We also briefly discuss molecular biology considerations, choice of sequencing platform, and the data analysis pipeline. Lastly, we discuss options for automating the pipeline and leveraging cloud-based platforms for data analysis.

### 2.1 parSEQ Process Description

parSEQ was developed to enable the efficient isolation of variants in a DNA pool into sequence-verified physically-separated containers that enable easy variant retrieval. We set up parSEQ to work with 384 well plates, with each well serving as a container throughput the process. The overall process is divided into five stages: clonal isolation, sequence barcoding, sequencing, data analysis, and finally cherry picking. DNA pools come in different formats (plasmid pools, phagemid pools, bacterial pools, yeast pools,…). We have chosen to work with bacterial pools as input to the parSEQ pipeline because of their robustness and ease of manipulation. Thus, if the DNA pool exists in some other format, it has to be subcloned and/or transformed into a bacterial pool. Any bacterial plasmid vector should work with parSEQ, and we have used multiple including pET12, pET19, and pET21. Clonal isolation aims at isolating individual bacteria from a bacterial pool into separate clonal wells. Sequence barcoding aims at indexing each well with well and plate specific barcodes. This is accomplished in two steps. The first step is a Taq polymerase colony PCR on each clonal well using 384 well-indexed primer pairs. The index of each primer pair points to one well of a 384 well plate. Post colony PCR, in the second step, each plate is pooled into a tube, and the tube is library prepped for sequencing. Within the library preparation process, we use indexed sequencing adapters that point to one of the 384 well plates. After library prep, all library-prepped plate samples are pooled together and sequenced. We used both Illumina MiSeq and ONT’s MinIon as sequencing readouts (library preps were performed accordingly). Post sequencing, data analysis on the raw sequencing output recovers the sequence contained in each well of the run. At this stage, a researcher can check all the unique sequences reported by parSEQ and then proceed to cherry pick the variants of interest.

### 2.2 Easy Implementation of parSEQ across Labs of Varying Scales

Implementing the parSEQ pipeline is technically feasible and economically viable, catering to labs of varying scales both in academia and industry. parSEQ can be implemented in different configurations depending on throughput needs and capital cost limitations. At one extreme end, manual clone picking, multi-channel pipettes liquid handling, outsourced NGS sequencing to low-cost providers, and desktop-based NGS analysis can be set up with minimal capital costs and expertise, at the expense of lower throughput sampling and higher sampled clone cost. This setup is similar to the setup used by evSeq ^14^. At the other end, automated clone picking, automated liquid handling, high-capacity in-house sequencers (e.g. ONT PromethION), and cloud-based analytics can provide standardization, very-high throughput sampling, and lower sampled clone cost. Multiple different configurations exist between these two extremes, such as using lower-cost & slower colony pickers, semi-automated liquid handling, and lower capacity in-house sequencing (e.g. ONT MinION).

### Necessary Equipment

- Clonal Isolation Setup. Multiple options are available:

**–** Bacterial cell sorter or FACS, available on most university campuses.
**–** Specialized instruments for single bacterial cell dispensing such as Cytena’s B.Sight or Namocell’s Hana.
**–** Microbial colony pickers such as molecular devices’ QPix series, Hudson Robotics’ RapidPick SP & RapidPick Lite Colony Pickers, and Singer Instruments’ PIXL colony picker.
**–** Manual Picking, if determination prevails! Numerous labs manually pick thousands of colonies routinely.
- Liquid Handling Solution

**–** Liquid Handling Robots such as Hamilton Starlet or any alternative liquid handler that incorporates open-source software for programming, negating dependency on proprietary protocol development, which can be both time-consuming and expensive. Typically, such solutions necessitate some level of expertise for establishing protocols and workflows.
**–** Lower cost semi-automated solutions such as Integra’s ViaFlo or Assist series are economical, efficient, flexible, and easy to set up but do require more hands-on time from the operator to run the pipeline.
**–** Multi-channel micro-pipette. Classic.
- Thermal Cyclers

**–** 384-well-plate block if decided to work with 384 well plates.
**–** 96-well-plate block if decided to work with 96 well plates.
- Sequencing Platform. Most academic campuses provide access to sequencing platforms via core facilities. Still, for those lacking access, alternatives exist:

**–** Employing service providers like Genewiz, which offers economical DNA NGS services such as Amplicon-EZ .
**–** Opting for one of ONT’s MinION Starter Packs, with some starter packs being available for $ 1,000 (Mk1B) or $ 5,000 (Mk1C) capital cost, including flow cells, consumables, and an integrated computer for the Mk1C.
- General Lab Equipment

This adaptability in equipment and process makes parSEQ a versatile tool, accessible to a broad range of labs.

### 2.3 parSEQ Running Costs

The primary costs associated with running the parSEQ pipeline are derived from barcoding colony PCRs (including plate-level pooling and purification) and sequencing.

- Barcoding Colony PCRs: No High Fidelity polymerase is required for parSEQ. We recommend any variation of HotStart Taq polymerase. Barcoding cost per well including all reagents, plastic-ware, plate-level pooling and purification, but excluding labor cost, will fluctuate around 20 cents depending on choice of consumables. In any case, this cost should not exceed 25 cents per well.
- Sequencing: Depending on the sequencing platform chosen, costs will vary.

**–** Employing a budget-friendly NGS service like Amplicon EZ will render a sequencing cost of approxi-mately $60 per plate, equating to less than 20 cents per well.
**–** Opting for ONT’s MinIon sequencing will result in an approximate cost of $50 per plate, translating to less than 15 cents per well. A noteworthy advantage here is the immediacy of results, contrasting with the approximate 10-day lead time of sequencing providers.

When incorporating ONT sequencing in-house, the comprehensive cost of parSEQ, including labor costs, is projected to not surpass 50 cents per well sampled. This denotes an over tenfold reduction in the per-well sequencing expense in comparison to conventional Sanger sequencing. This substantial reduction in the cost per variant sampled enables the adoption of a sequence-first-screen-later approach.

### 2.4 parSEQ Timelines

When contrasted with workflows incorporating Sanger sequencing, parSEQ maintains analogous timelines. The entire pipeline, capable of sequencing up to 20-24 plates, can be executed within a single week using our automated setup and four 384-well plate thermal cycling blocks. In Figure 3, we juxtapose the timelines associated with deploying parSEQ against those of Sanger sequencing, focusing on the output of a panning procedure for 16x 384-well plates of selected colonies.

**Figure 3:**
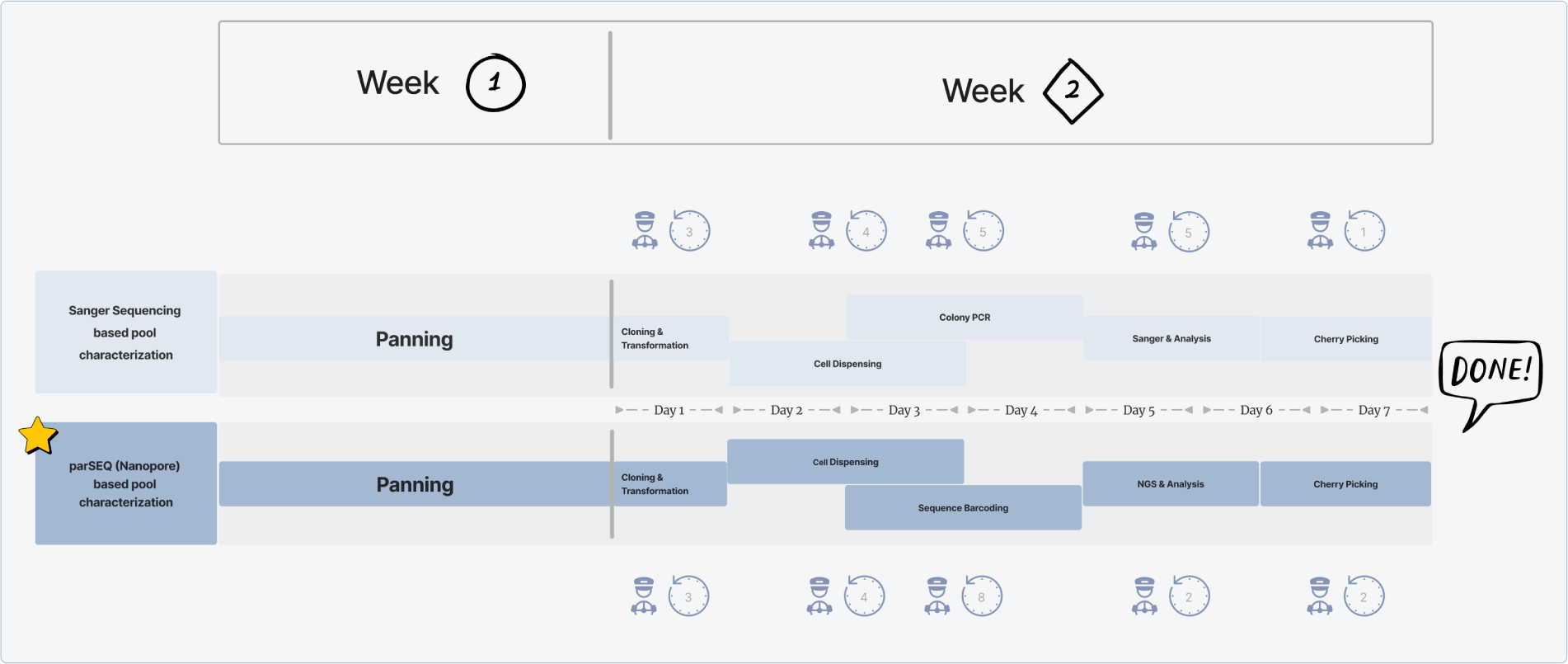
Timeline comparison between parSEQ and Sanger sequencing workflows in the context of protein binder discovery. The process begins when a researcher creates an enriched library of potential binders using a display method and successive rounds of selection against a target antigen (panning). The next step is to characterize the binders in this enriched library. Using parSEQ, characterizing a sizable sample of 6,144 clones (across 16x 384-well plates) can be completed in approximately 7 days. This is on par with the timeframe it would take to process the same number of samples with Sanger sequencing, given that the fundamental molecular biology steps are similar in both approaches. However, where parSEQ really stands out is in cost efficiency – it’s about ten times cheaper than the Sanger method. This significant cost reduction makes it viable to recover unique variants directly from display pool outputs, a strategy too expensive when using Sanger sequencing. Thus, parSEQ introduces a cost-effective alternative that makes comprehensive library screening accessible and practical for researchers.

### 2.5 Molecular Biology and Sequencing Platform Considerations

#### 2.5.1 Molecular Biology Considerations Barcoding Colony PCR

- Choice of polymerase:

There’s no need for costly high-fidelity polymerases in the barcoding colony PCRs. **Standard Taq polymerases** are adequate. Any errors introduced during polymerase amplification (as well as those from sequencing) will be randomly scattered across the DNA strands that have been amplified or sequenced. Based on our calculations and experience, these random errors are overshadowed by the underlying sequence data, provided a sufficient number of reads per well are analyzed. Specifically, around 50 reads are required when sequencing with Illumina, while approximately 300 are necessary when sequencing with ONT v14 chemistry and using Dorado’s super accuracy base calling models. We also recommend using **Hot-Start Taq polymerases**.This approach facilitates the simultaneous preparation of all barcoding PCR plates, which can then be stored and sequentially placed on thermal cycling blocks. In our operations, for instance, it takes about 4 minutes to prepare a single barcoding PCR plate, whereas the actual PCR reaction requires roughly 2 hours on a thermal cycling block. Using our automated setup, we can prepare 16 PCR plates in less than 1.5 hours, while running these plates on the four available thermocycling blocks needs 8 hrs approximately. Consequently, the duration of the PCR reaction emerges as the primary bottleneck in our pipeline, and likely yours as well. Therefore, using Hot-Start Taq is practical since it permits batch preparation of PCR plates, which can be stored as the PCR reactions unfold.

- Barcoding oligo design:

We employed nxCode by the Hannon Lab to design 384 unique nucleotide barcodes, each 9 bases long, assigning one barcode to each well in a 384-well plate. These barcodes are expected to have a decoding accuracy of 0.997 with Illumina sequencing. Following this, we connected each barcode to two separate universal buffer regions; one tailored for forward primers and the other for reverse primers. Consequently, every well was equipped with a primer pair consisting of one forward and one reverse primer. Although the forward and reverse barcoding primers in each well contain the same barcode, they are linked to different buffer zones. These universal buffer areas are designed to align with corresponding zones on the set of template amplifying primers (refer to Figure 4).

- PCR reaction setup and thermocycling protocol:

**Figure 4:**
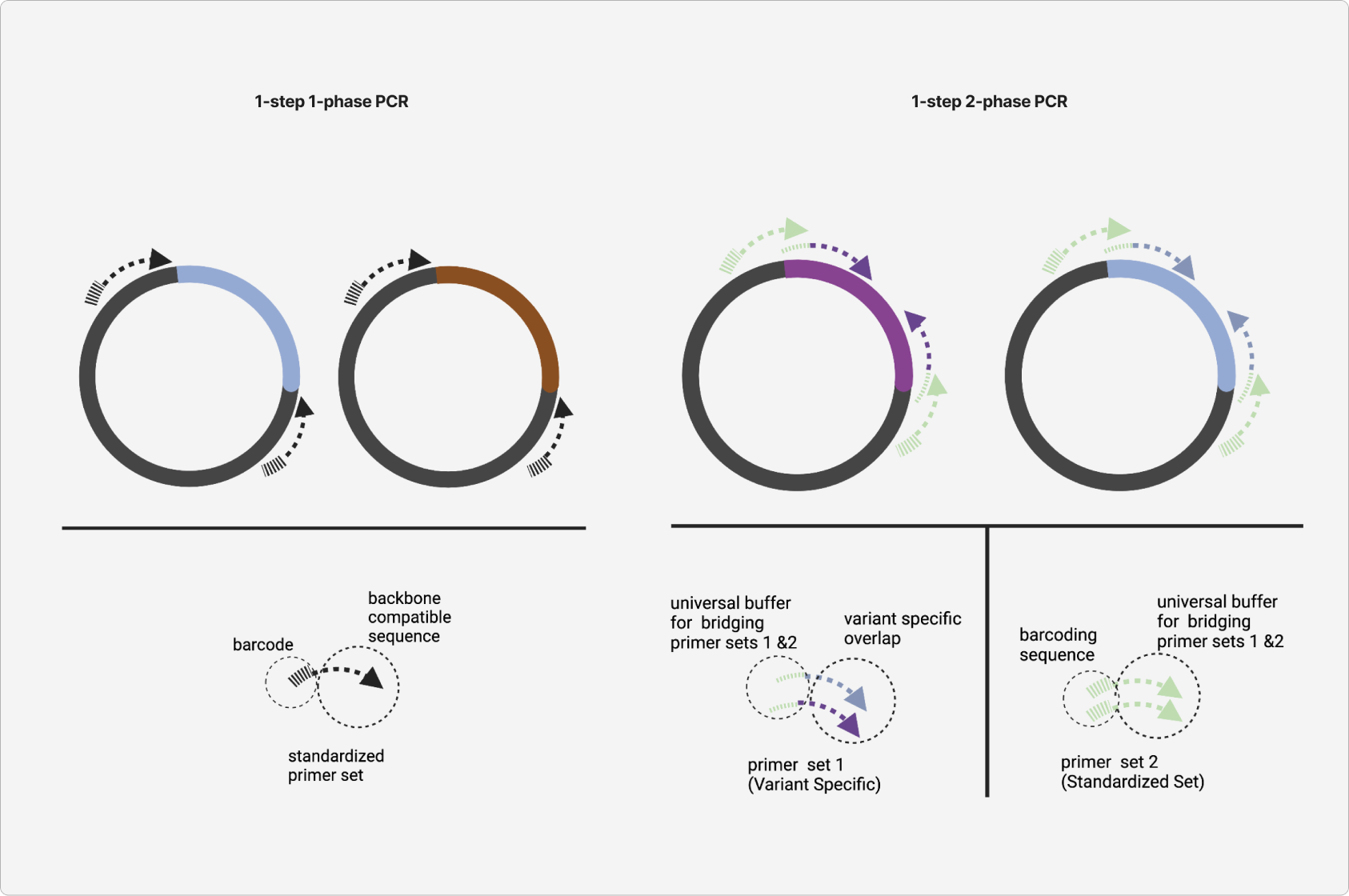
Advantages of ONT’s unlimited sequencing length for standardizing components in amplicon sequencing. ONT’s sequencing technology allows for sequencing without length restriction, from as short as 20 base pairs to as long as millions of base pairs (megabasepairs). This feature is beneficial for techniques like parSEQ’s pooled amplicon sequencing. With ONT, one can use a single set of barcoded primer that anneals to a common sequence in the plasmid backbone to simultaneously amplify and barcode any DNA segment cloned into the plasmid. This approach is not possible with length-limited sequencing platforms like Illumina, because the sequencing read won’t cover the full length of the longer amplicant. . ONT’s flexibility enables us to use one 384 set of barcoded oligonucleotides, allowing for 1-step, 1-phase barcoding PCRs. This approach is simpler compared to the 1-step, 2-phase barcoding PCRs required for length-restricted platforms like Illumina. Typically, our parSEQ workflow involves 384-well plates, necessitating 384 unique indexed oligo duplexes. With ONT, we can design oligos that anneal to a common plasmid region, meaning that we can create 384 oligo duplexes, each consisting of a backbone-binding sequence at the 3’ end and a barcode at the 5’ end. This uniform approach isn’t feasible with length-restricted methods like Illumina, forcing us to engage in 1-step, 2-phase PCRs (with all reagents added at once, but involving two phases: the first for inner primer amplification, and the second for external barcoded primer amplification). This approach hinders standardization and reduces PCR product quality. Also note that in cases of long variants, Illumina’s sequencing length limitation makes it difficult to cover the entire variant length using a single amplicant, and necessitate multiple amplifications per well. This is quite impractical, and leads to a lower quality of sequencing readout. Note that the 1-step 2-phase PCR reactions were used in the case studies presented in this technical report.

The standard PCR reaction setup used for experiments in this report is a 10 µl single-step, two-phase reaction (Figure 4, right). We introduce the first set of primers (also known as bridging or template-amplifying primers) into the reaction at a final concentration of 50 nM, and the second set (the barcoding primers) at 500 nM. These primer sets are designed so that the first set has a significantly higher melting temperature than the second. This distinction is important because, in the first phase of the reaction, a higher melting temperature is utilized for the initial 8 cycles, enabling only the first set of primers to amplify the template plasmid. We anticipate that nearly all the primers from the first set will be used up after these 8 cycles. During the second phase, the melting temperature is lowered, allowing the second set of primers to initiate amplification over 25 cycles, thereby amplifying and applying barcodes to the products of the first-phase amplification. Liquid Bacterial culture samples are introduced to the reaction at a volume of 0.2 µl, either directly or via low-volume stamping. The PCR protocol starts with a 1.5-minute step at 95°C to lyse bacterial cells. The PCR reaction conditions can be summarized as follows:

**–** Reaction volume: 10 µl

**–** Primer set 1 concentration: 50 nM

**–** Primer set 2 concentration: 500 nM

**–** Phase 1: 8 cycles -> engage primer set 1

**–** Phase 2: 25 cycles-> engage primer set 2

**–** Phase 1 melting Temp = Phase 2 melting Temp + 8-10°C

**–** Bacterial template samples are added at a volume of 0.2 µl (lower volumes give better results)

**–** Initial 90 sec bacterial lysis step at 95°C

This PCR reaction setup is compatible with workflows involving both Illumina and ONT as a sequencing platform. Having two phases of amplification with internal primers serving as the template amplification primers, and external primers serving as barcoding primers, allows us to separate template amplification from barcoding . Note that primer sets 1 and 2 bridge over each other to allow this 2 phase amplification/barcoding (Figure 4). A key advantage is the adaptability provided by the set of 384 barcoding primers; they can be consistently applied across various projects with different templates by merely switching the primers used in the first step (a set of two primers). This adaptability is vital for Illumina sequencing, where the readout length is constrained by the kit cycle number, making it essential to have flexibility in selecting primer annealing locations. For ONT sequencing, this flexibility is less crucial. The reasons for this discrepancy are discussed in Figure 4.

### Plate pooling

After the PCR process, EDTA is individually added to each well at a final concentration of 40 mM before the samples are combined. This step is crucial as the EDTA halts polymerase activity before pooling, preventing any recombination between variants from different wells. The contents of each plate are subsequently pooled into one tube.

### DNA sample purification

After pooling the samples, and also during subsequent stages of the library preparation protocol, sample purifications are carried out using Beckman’s AMPure XP Beads (Product No. A63881), according to manufacturer protocols.

### DNA concentration measurements

DNA concentration measurements at various points post-pooling are performed using Qubit dsDNA Quantification Assay Kits (Product No. Q32851).

#### 2.5.2 Choice of Sequencing Platform

In its initial configuration, parSEQ relied on Illumina MiSeq for sequencing, as the accuracy of ONT sequencing at the time wasn’t sufficient for reliable identification of well consensus. However, advancements in ONT’s V14 chemistry, combined with the R10.4.1 pore, and enhanced by Dorado’s high-precision basecalling models (SUP, v4.2.0), have significantly improved accuracy. This progress ^15,16^ established ONT’s MinIon as a dependable option for sequencing in parSEQ.

We have since transitioned to using the ONT MinIon Mk1C platform as our preferred method for parSEQ sequencing, owing to several advantageous features:

- Sequencing adaptability to any read length, ensuring streamlined operation. In figure Figure 4, we detail how the unrestricted sequence length of ONT sequencing enables standardizing amplicon sequencing for parSEQ. Standardization is important because it translates to lower cost, faster turnaround times, fewer errors, and higher pipeline efficiency.
- Enhanced run flexibility, including chip reuse.
- Minimal upfront cost, offering setup flexibility.
- Reduced cost of sequencing.
- Availability of real-time sequencing readout.

#### 2.5.3 Sequencing Library Preparation

The preparation of sequencing libraries allows for the pooling of samples from different plates into one sequencing run by utilizing indexed sequencing adapters (plate-level library multiplexing).To mitigate the risk of DNA variant recombination during PCR amplification, we employ PCR-free library preparation kits for both Illumina and ONT platforms. These kits utilize ligation-based workflows to attach indexed NGS adapters to our samples.

For Illumina sequencing, we employ the IDT xGEN™ DNA MC UNI Library Prep Kit. In the case of ONT sequencing, we use ONT’s Native Barcoding Kit V14 (SQK-NBD114). Both of these kits provide the option to extend adapters with up to 96 unique indices, facilitating the multiplexing process for up to 96 plates in one run. In practice, we have seen that runs should not multiplex more than 24-28 plates when using the ONT Minion flow cell. For multiplexing 96 plates in one run, using ONT’s PromethION2 or GridION systems is more appropriate.

### 2.6 parSEQ Data Analysis

We employ a Python-based pipeline to analyze parSEQ NGS results. This pipeline integrates multiple open-source libraries and packages, with the main goal of identifying the specific DNA sequence in each well that parSEQ investigates.

#### Preparing and Filtering Raw Data Files

- For basecalling on the ONT MinIon raw data, we use the SUP, v4.2.0 Super accuracy basecalling model with Dorado v0.2.5B. It’s important to note that Dorado necessitates GPU acceleration. In our setup, we employ a custom AWS EC2 instance equipped with an Nvidia T4. Additionally, Dorado generates only BAM files, so samtools are necessary to convert these to fastq. If you lack access to GPU acceleration, Dorado can run on Apple’s silicon chips (M1/2 series).
- For Illumina based sequencing, standard basecalling on the MiSeq is used.
- We use Fastp v0.20.1^17^ to filter raw reads by quality and length. Fastp is also used for merging paired-end reads when sequencing is done with Illumina.

#### Demultiplexing

- Plate level indexing is brought by indexed library prep sequencing adapters used for both Illumina and ONT sequencing. Plate-level demultiplexing is thus performed using standard Illumina MiSeq barcode demultiplexing options in case of Illumina sequencing. For ONT, plate level demultiplexing can be performed using Guppy.
- Freebarcodes v3.1.0^18^ is used for well-level demultiplexing of the reads, allowing a search error of 1 during the Freebarcodes demultiplexing process.
- After demultiplexing with Freebarcodes, we employ the edlib v1.3.9 library ^19^ and the Biopython v1.81 package ^20^ to detect forward and reverse reads, and to match reads to the appropriate wells.

#### Alignment

- For multi-sequence alignments at the well level, we utilise either MUSCLE v3.8^21^ or MAFFT v7.490^22^.

#### Well-level Consensus

- To establish the consensus sequence at the well level, we use the Bio.Align.AlignInfo module from Biopython. A consensus threshold of 0.8 is used for ONT-based sequencing, and a consensus threshold of 0.9 is used for Illumina based sequencing.

### 2.7 Pipeline Automation

To enable parSEQ to scale efficiently with a high pool sampling rate, automation of the wet-lab process is key. As detailed previously, parSEQ relies on a series of molecular biology techniques to retrieve DNA from isolated clones and prepare it for sequencing. These processes are particularly amenable to automation as their core requirement is liquid handling, for which many commercial solutions exist.

Automated liquid handling solutions have traditionally been leveraged by large companies and well-funded labs as they were prohibitively expensive. However, lower cost options have recently made it to market (e.g. Opentrons). Moreover, even semi-automated solutions such as Integra’s ViaFlo product line can massively improve the throughput of routine and repetitive experiments, similar to the ones required by parSEQ.

We chose the Hamilton STARlet, a modular system, to fully automate the parSEQ process. The system’s 384-channel head is ideal for handling 384-well plates, enhancing our throughput. Initially, we had concerns about the closed nature of its VENUS control software, which required support from Hamilton and resulted in prolonged development timelines. To overcome this, we turned to PyHamilton ^23^, an open-source Python framework created by researchers at the MIT Media Lab for programming Hamilton liquid-handling robots. PyHamilton enables researchers to develop automated liquid-handling protocols for Hamilton robots using Python. This integration means that established software strategies, including exception handling, version control, and object-oriented programming, can be applied directly to protocol development. With PyHamilton, we experienced a significant acceleration in our protocol development, shortening timelines notably. We developed our parSEQ-PyHamilton library on top of PyHamilton to automate the parSEQ pipeline and maximize walk-away time for operators. An illustration of our parSEQ-PyHamilton automation stack is provided in Figure 5.

**Figure 5:**
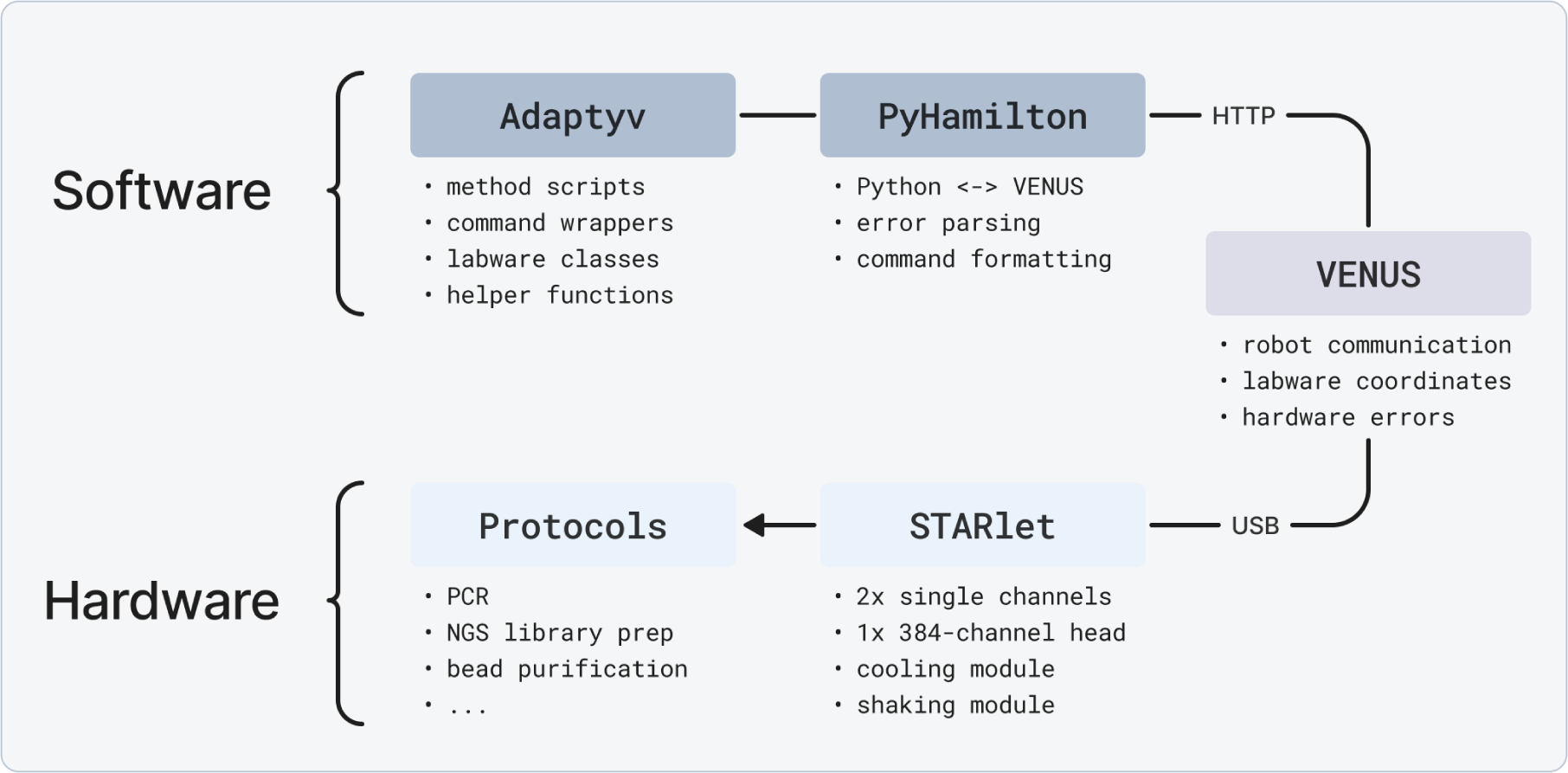
Scheme of the parSEQ automation stack built using Adaptyv-PyHamilton and Hamilton STARlet robot. PyHamilton acts as the communication layer between our protocols and Venus (proprietary Hamilton control software) via an HTTP server. In turn, Venus processes these user commands packets into actual commands executable by the robot. Our higher level parSEQ library codes protocols by integrating lower level command wrappers for simple robot actions such as pipetting steps and robot movements.

Our parSEQ-PyHamilton library builds on the PyHamilton framework and adds several new features, including state management for error recovery and tracking of each step in a protocol. It also adds commands for using a 384-channel pipetting head, extends support for common labware formats, and implements protocols as Python scripts callable from a main CLI script. Additionally, the library provides wrappers to simplify common robot movements, labware classes that implement physical labware as pandas dataframes, and notifications via Slack API when user intervention is required.

To better demonstrate the gains brought by our automation stack, two example protocols are detailed below:

1. **Pooling Protocol:** This protocol takes a post-PCR 384-well plate as input and outputs a single 2 mL Eppendorf tube containing an equal amount of liquid from each well. The process involves adding EDTA to all 384 wells to inhibit the PCR reaction. The protocol can handle up to 24 plates in one run without any user intervention, and takes around 10 minutes per plate. This is a significant improvement compared to the typical 30 minutes required when performed manually. We’ve compared the time and resources needed for three different approaches of conducting this protocol — fully automated (with our system), semi-automated (using Integra’s Assist platform), and manual. You can see the details in Table 1.
2. **Barcoding Colony PCR Protocol:** This procedure requires 384-well plates filled with bacterial liquid cultures, a master mix containing Taq polymerase (in a single well plate), and a single 384-well plate loaded with barcoding primer pairs. The end result is an equivalent number of PCR ready 384-well plates that reflect the original layout of the bacterial cultures. These plates are now ready for the thermal cycling process. We’ve compared the time and resources needed for three different approaches of conducting this protocol — fully automated (with our system), semi-automated (using Integra’s Assist platform), and manual. You can see the details in Table 2.

**Table 1:** Operational Gains from Automating the Pooling Protocol.

|  | **Manual** | **Semi-automated** | **Automated** |
| --- | --- | --- | --- |
| **Time** | 30 min/plate | 17 min/plate | 11 min/plate |
| **Consumables (per plate)** | 12x P20 tips<br>1x P1000 tips<br>1x 96-microplate | 104x P20 tips<br>1x P1000 tips<br>1x 96-microplate | 104x P50-tips<br>2x P300 tips<br>1x 96-microplate |
| **Consumables (per run)** | 12x P20 tips<br>1x reservoir | 384x P20 tips<br>1x reservoir | 384x P50 tips<br>1x reservoir |
| **Intervention** | required | required | none |

**Table 2:** Operational Gains from Automating the barcoding colony PCR.

|  | **Manual** | **Semi-automated** | **Automated** |
| --- | --- | --- | --- |
| **Time** | 24 min/plate | 9 min/plate | 4 min/plate |
| **Consumables (per plate)** | 768x P20 tips<br>1x 384-microplate | 768x P20 tips<br>1x 384-microplate | 768x P50 tips<br>1x 384-microplate |
| **Consumables (per run)** | 12x P20 tips<br>1x reservoir | 384x P20 tips<br>1x reservoir | 384x P50 tips<br>1x reservoir |
| **Intervention** | required | required | none |

Developing new automated methods for synthetic biology processes is straightforward using our library and templates. We (biologists) naturally think about an experimental method in a step-wise fashion: aspirate this, move here, dispense while mixing, discard some used labware, etc. parSEQ-PyHamilton keeps things simple and familiar: each protocol is written with this logic in mind, allowing you to easily add, remove or reorder steps as the process dictates.

In general a new protocol can be developed in a few steps:

1. Define layout and labware in VENUS
2. Define main steps of the protocol
3. Breakdown repetitive steps in for loops
4. Define start and stop conditions
5. Use existing protocols and command wrappers to build your script

Finally, it’s worth noting that there are some limitations to our parSEQ-PyHamilton library. It relies on VENUS, Hamilton’s proprietary software, to create layout files, and users need to have basic Python knowledge. However, the potential increase in throughput (essential for parSEQ) is well worth these hurdles.

Overall, our parSEQ-PyHamilton library, built on top of the PyHamilton library, has enabled us to efficiently develop our automation protocols, saving time and increasing throughput. By leveraging the open-source PyHamilton module, we’ve created a library of protocols that can be easily shared and adapted to suit your needs. In parallel we have developed an automated cloud pipeline to process parSEQ data, which we will detail in the next section.

### 2.8 Implementation of Analysis on the Cloud

#### 2.8.1 Why Implement Analysis on the Cloud?

Early on, we were processing and analyzing our parSEQ NGS data using Jupyter notebooks and Python scripts on Amazon Web Services (AWS) EC2 instances. This method was convenient and relatively simple to setup. The flexibility of Python scripts and Jupyter notebooks made setting up and changing scripts easy, plus they integrated well with existing bioinformatics tools and packages.

However, as our operations scaled up and we began processing large volumes of plates regularly, we faced a challenge. Data began accumulating faster than we could analyze it, and the standalone EC2 instances we were using couldn’t keep up with the workload. Furthermore, we struggled to keep track of the relationships between different sets of data — it was hard to know which scripts had generated which results and it was cumbersome to link raw and intermediate data with the final stored data. This lack of organization made it difficult to analyze and compare results across different runs.

#### 2.8.2 Requirements for Analysis on the Cloud

To address computational delays and improve data tracking, we established a set of criteria for our analysis pipeline, detailed below.

First, we anticipated handling numerous projects, each consisting of tens of 384 well-plates, with expectations for this volume to grow. The most demanding task, computationally, was the multi-sequence alignments conducted on each well. With up to 384 sequence alignments necessary per plate, our pipeline needed sufficient scalability to perform these tasks in parallel within a reasonable time frame. This capability would enable, in theory, a set of 12 plates (encompassing as many as 4,608 sequence alignments) to be processed in the same duration needed for a single well’s multi-sequence alignment, significantly speeding up the operation (Figure 6).

**Figure 6:**
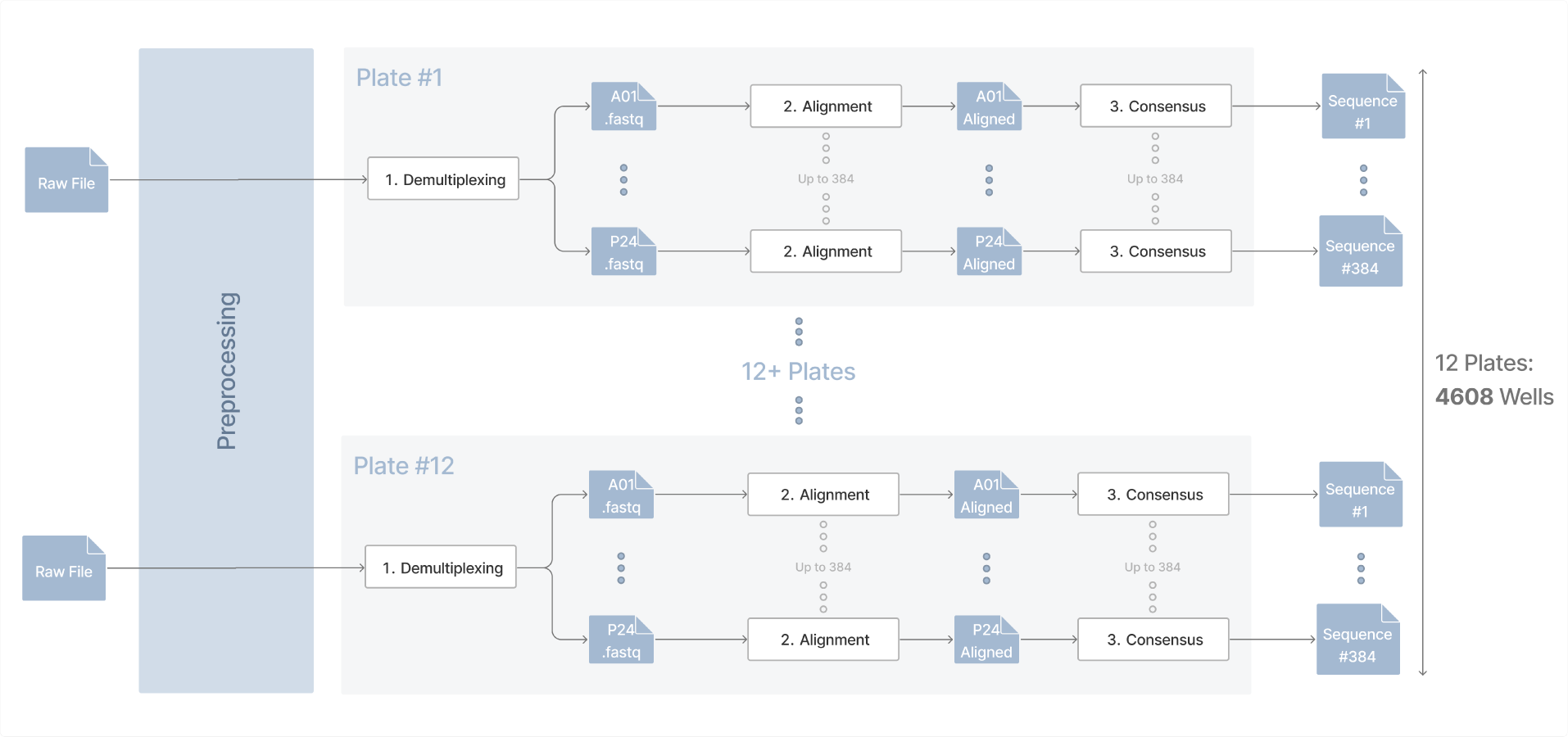
Outline of the main requirements for implementing parSEQ analysis. Raw sequencing files corresponding to different plates undergo almost identical analysis to recover well-level consensus sequences. The main steps in this analysis are raw data processing (or preprocessing), demultiplexing, alignment, and consensus. Pre-processing will vary based on the sequencing platform (Illumina or ONT) used, and the basecalling method used for ONT. Remaining steps are identical.

**Figure 7:**
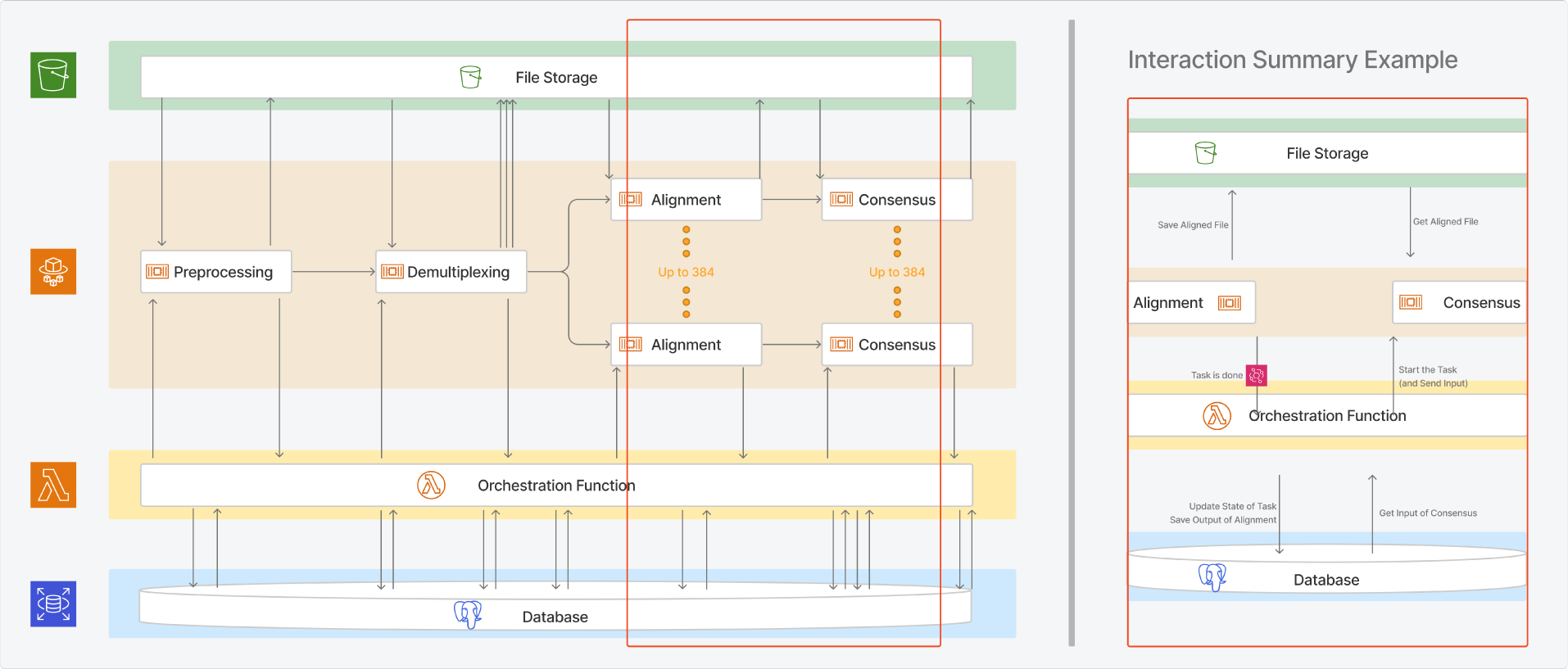
Cloud analysis pipeline. We built our analysis pipeline using Amazon Web Services infrastructure. S3 is used for file storage, an SQL Postgres AWS RDS database is used for structured data storage, lambda functions are used for tasks needing minimal resources but high parallelization, and the ECS Task service is used for resource-intensive tasks -such as demultiplexing and alignment-that utilize docker images stored in AWS’s ECR. A central lambda function manages the orchestration of tasks between the database and ECS task service.

We also recognized that every parSEQ run generates substantial data, which various teams might need to access, including those working on synthetic biology as well as machine learning. Consequently, we required a scalable storage solution accommodating diverse file formats (like fasta, fastq, bam, etc.), as well as final structured tabular data.

Crucially, we also needed a system for connecting this data, not just to each other, but also to the specific processes or tasks that generated it. This involved creating a method whereby any file or data piece could be traced back through each step that led to its creation, and similarly, each process or task could be traced forward to every piece of data it produced.

In addition, our workflow often involves updates and alterations to the code that runs specific processes, whether for optimization, algorithm enhancement, or other reasons. We wanted these updates to be integrated into the cloud-based tasks seamlessly, automatically adjusting whatever infrastructure components were necessary. This adaptability serves a dual purpose: it allows us to select specific code versions for tasks and ensures that our data analysis can always indicate which code version produced the final results.

Furthermore, our aim was not just to modify existing tasks but to have the flexibility to introduce new tasks, substitute old ones, rearrange their sequence, and reuse them across different projects. Since some tasks are applicable to multiple pipelines, a modular approach was preferable. This way, as we develop more tasks, we gain additional, interchangeable components for future pipelines.

Looking ahead, we envisioned an infrastructure capable of extending even to laboratory tasks. This comprehensive approach is crucial because pipeline analysis and optimization aren’t confined to computational tasks. For the fullest understanding and enhancement of our end product’s production and quality, we need insights into the lab processes as well. Therefore, we intended our system to be readily expandable, with future updates able to incorporate and document this lab information from inception to the conclusion of the pipeline.

#### 2.8.3 Analysis Pipeline

Let’s illustrate our approach with an example: if we decide to switch from ONT to Illumina sequencing, we’d need different data prep for Illumina’s raw data output before demultiplexing. We’d write the code for this step, package it as a docker container, upload it to the elastic container registry (ECR), and set an elastic container service (ECS) task definition pointing to this new image. This strategy enhances modularity and parallelization.

With these tasks as our building blocks, we needed a system to coordinate them—knowing when to initiate the next task upon the completion of a previous one. We chose to create our orchestration function as a lambda function, triggered by any pipeline-related event, using AWS Event Bridge to report ECS task events to this function. It decides whether to initiate subsequent tasks or halt the pipeline based on these events. If we add a task (like the pre-demultiplexing paired-end raw fastq file merge for Illumina), we only need to adjust the orchestration function’s code to incorporate this new step.

This structure gives us a flexible pipeline where tasks can be added, removed, or reordered. To track tasks, inputs, and outputs, we implemented two storage forms: an SQL database (Postgres on AWS RDS) for structured data and AWS S3 for file storage. The Postgres database, our “single source of truth,” contains references to all files in S3 and also plays a role in orchestration and system state management. It records every task interaction with the files, the parameters used, and other task-specific information. This database is accessible to all pipeline components, particularly the orchestration function, ensuring up-to-date information on the system’s status is available for queries at any time.

We also aim to track the version of each task producing specific outputs, and potentially run different task versions concurrently. For instance, we may have various demultiplexing algorithms for different projects. Since we use GitLab for version control, we integrated GitLab’s Continuous Integration (CI) services. When new code is committed, a new docker image is automatically built, added to ECR, and a corresponding ECS task definition is created. This information is logged in our SQL database, allowing us to specify task versions for each pipeline run and later determine the exact code version used for each output during data analysis.

Our database’s design is process-agnostic—it can support any data-generating process. It can also potentially accommodate details of lab procedures like PCR, recording all pertinent information and parameters. This system’s extensibility is broad, and caters to various needs.

#### 2.8.4 Stats and Visualizations

The core pipeline, previously discussed, has a parallel section dedicated to using data for statistics and visualizations. For example, after demultiplexing, a heatmap of read counts for each plate is generated simultaneously with other tasks like alignment. This heatmap, once completed, is saved and logged in the database, with the file stored in S3. For other stats and visuals which aggregate data from multiple consensus sequences across various plates or an entire run, the orchestration function monitors progress and initiates the plot’s creation once all necessary data is available. An example is a bar plot of average sequence lengths across a run, initiated only after gathering all consensus sequences across a run. Adding or removing these plot-generating tasks is as straightforward as it is for main tasks like demultiplexing. The system’s features, including code tracking, data storage, and orchestration, apply to these tasks as well. This approach maintains consistency and efficiency in both primary data processing and auxiliary functions like statistics and visualization. In addition, all data collected during all runs is readily available for querying and additional visualization.

## 3 Case Studies

We present the following case studies to demonstrate how parSEQ can be used to maximise the capture of sequence-function data-sets in different contexts. Case study 1 showcases how parSEQ can be used to facilitate high-throughput variant retrieval from a DNA pool. Case study 2 demonstrates parSEQ’s ability to retrieve expression ready bacterial clones from bacterial pools. Case study 3 illustrates how parSEQ can be used to source DNA for testing denovo designed proteins. Lastly, we propose case study 4. Case study 4 explores the potential use of parSEQ in preparing sequence-verified, clonally isolated targeted mutational libraries. As we haven’t conducted experiments to validate case study 4, it could be an area for future users to explore.

### 3.1 Case Study 1: parSEQ facilitates a deep exploration of display pool outputs and retrieves unique sequences with minimal enrichment

We obtained the phage display output pool of a nanobody library panned against an antigen from a client. We first ran NGS to elucidate its enrichment profile. NGS was run using ONT’s Minion, kit V14 chemistry, and flow cell R10.4.1. Basecalling was executed using Dorado’s super accuracy models (SUP, v4.2.0). Subsequent to basecalling, fastp managed the read filtering for quality and length. The filtered reads were then clustered using CD-HIT ^24,25^ with sequence similarity thresholds ranging from 0.8 to 0.97 in increments of 0.01. Average inter/intra cluster distances at each threshold were calculated and plotted against the threshold value (Figure 8,a). A pivotal elbow point emerged at a similarity threshold of 0.92, dictating the basis for ensuing analyses. Post clustering at sequence similarity threshold of 0.92, the majority of the 4952 clusters, predominantly containing single-digit sequences, were pruned, retaining approximately 400 clusters (Figure 8,b). A sampling simulation on this refined read distribution (Figure 8,c), predicted the retrieval of approximately 280 unique sequences from 3,840 (10x 384 well plates) samples under optimal conditions. Considering an 85% efficiency in our parSEQ pipeline (ratio of the number of wells sequences retrieved to the number of bacterial culture wells processed), we projected a retrieval of around 238 unique sequences.

**Figure 8:**
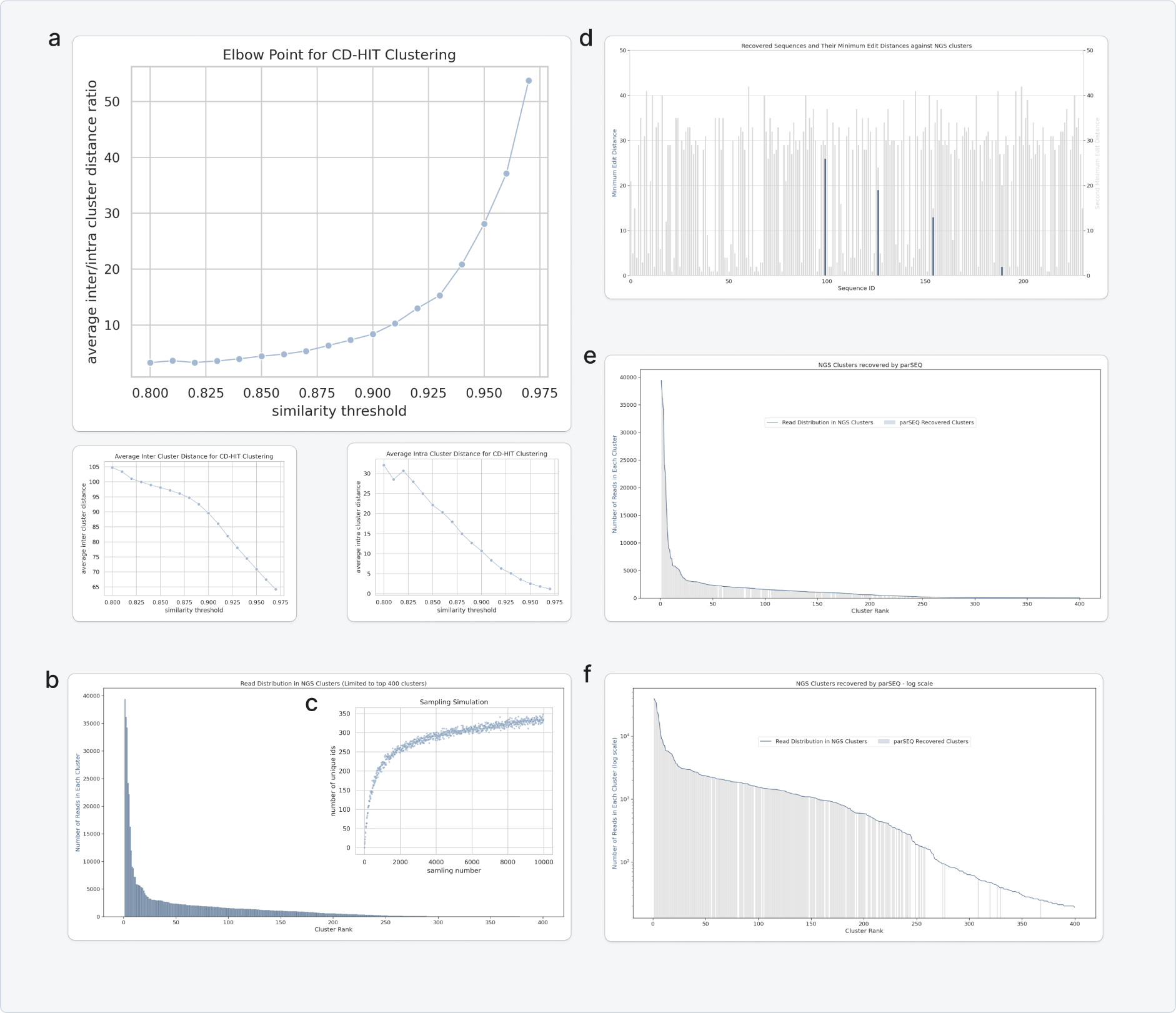
Case Study 1:parSEQ enables efficient variant retrieval from DNA pools. Comparison of clustering on NGS reads with parSEQ retrieved sequences. We obtained the phage display output pool of a nanobody library panned against an antigen from a client. We first ran NGS to elucidate its enrichment profile. NGS was run using ONT’s Minion, kit V14 chemistry, and flow cell R10.4.1. Basecalling was executed using Dorado’s super accuracy models (SUP, v4.2.0). Subsequent to basecalling, fastp managed the read filtering for quality and length. **a)** The filtered reads were then clustered using CD-HIT with sequence similarity thresholds ranging from 0.8 to 0.97 in increments of 0.01. Average inter/intra cluster distances at each threshold were calculated and plotted against the threshold value. A pivotal elbow point emerged at a similarity threshold of 0.92, dictating the basis for ensuing analyses. **b)** Post clustering at sequence similarity threshold of 0.92, the majority of the 4952 clusters, predominantly containing single-digit sequences, were pruned, retaining approximately 400 clusters. **c)** A sampling simulation on this refined read distribution, predicted the retrieval of approximately 280 unique sequences from 3,840 (10x 384 well plates) samples under optimal conditions. Considering an 85% efficiency in our parSEQ pipeline (ratio of the number of wells sequences retrieved to the number of bacterial culture wells processed), we projected a retrieval of around 238 unique sequences for the said sampling volume. **d)** We ran parSEQ on 10x 384-well plates sampled from the pool, employing ONT’s Minion with kit 14 chemistry and flow cell R10.4.1 for sequencing readout. Basecalling was performed using Dorado’s super-accuracy basecalling models (SUP, v4.2.0), followed by read filtering for quality and length using fastp. Subsequently, freebarcodes software decoded the barcodes, allocating the sequencing reads to their respective wells. Mafft v7.490 aligned the reads in each well, enabling the recovery of a consensus sequence for each well. A comparative analysis of cross-well consensus sequences revealed 228 unique sequences, aligning with our sampling simulation. These sequences were then matched with the clusters’ consensus sequences, utilizing Levenshtein edit distance as the similarity index. For each unique sequence recovered, we identified the cluster with the least distance and the one with the second-least distance, and plotted the results. Except for four sequences, we located clusters with matching consensus sequences, exhibiting a Levenshtein distance of 0. We then compared the enrichment profiles of the recovered sequences in normal scale **(e)** and in logarithmic scale **(f)**, observing, as anticipated, a lower recovery of NGS clusters with lower pool enrichment values. The recovered sequences/clusters with the lowest enrichment had 25 reads out of the 579,860 total NGS reads processed, representing approximately 0.004% enrichment.

We ran parSEQ on ten 384-well plates sampled from the pool, employing ONT’s Minion with kit 14 chemistry and flow cell R10.4.1 for sequencing readout. Basecalling was performed using Dorado’s super-accuracy basecalling models (SUP, v4.2.0), followed by read filtering for quality and length using fastp.

Subsequently, freebarcodes software decoded the barcodes, allocating the sequencing reads to their respective wells. Mafft v7.490 aligned the reads in each well, enabling the recovery of a consensus sequence per well. A comparative analysis of cross-well consensus sequences revealed 228 unique sequences, aligning with our projections. A random selection of 10 of these retrieved sequences were confirmed using Sanger sequencing.

These parSEQ retrieved sequences were then compared to the clusters’ consensus sequences, utilizing Levenshtein edit distance as the similarity index. For each unique sequence recovered, we identified the cluster with the least distance and the one with the second-least distance (Figure 8,d). Except for four sequences, we located clusters with matching consensus sequences for all retrieved unique sequences, exhibiting a Levenshtein distance of 0.

We then compared the pool enrichment profiles of the recovered sequences (Figure 8,e in normal scale and Figure 8,f in logarithmic scale), observing, as anticipated, a lower recovery of NGS clusters with lower pool enrichment values. The recovered sequences/clusters with the lowest enrichment had 25 reads out of the 579,860 total NGS reads processed, representing approximately 0.004% enrichment. These findings highlight the strength of parSEQ in isolating unique sequences with minimal enrichments in DNA pools.

Having validated the parSEQ process, our objective shifted to contrasting the enrichment profiles between parSEQ and conventional NGS. For every distinct sequence retrieved via parSEQ, we recorded the number of parSEQ wells containing the said sequence, termed as hits or well support. Subsequently, for each unique sequence, we determined the count of NGS reads exhibiting the closest relation to this distinctive sequence, utilizing Levenshtein distances for matching NGS reads to unique parSEQ sequences. The enrichment profiles of unique variants given by parSEQ and NGS are juxtaposed in Figure 9,a.

**Figure 9:**
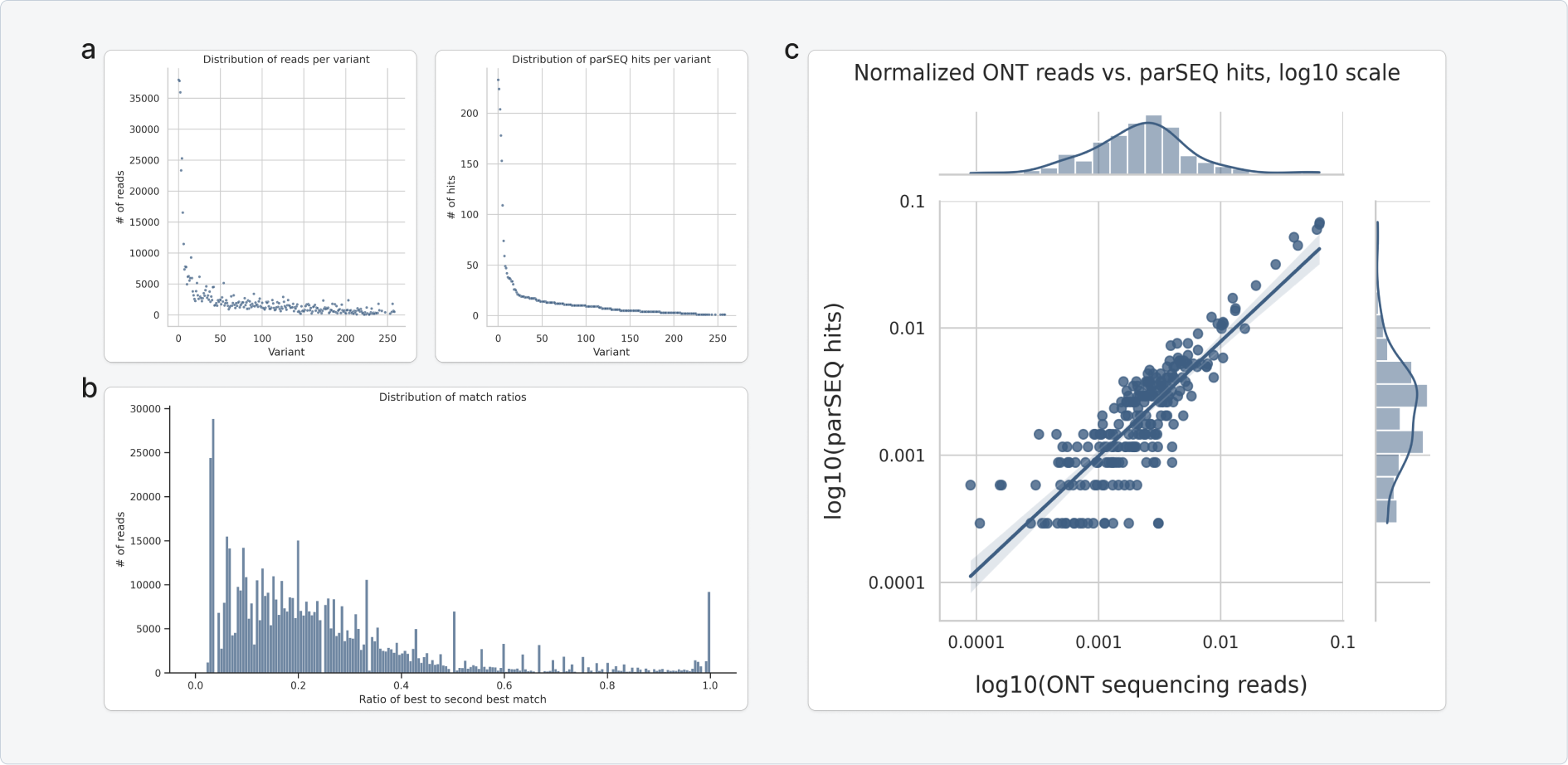
Case Study 1: parSEQ enables efficient variant retrieval from DNA pool. Aligning NGS reads to parSEQ retrieved variants. **a)** For every distinct sequence retrieved via parSEQ, we recorded the number of parSEQ wells containing the said sequence, termed as hits or well support. Subsequently, for each unique sequence, we determined the count of NGS reads exhibiting the closest relation to this distinctive sequence, utilizing Levenshtein distances for matching NGS reads to unique parSEQ sequences. The enrichment profiles of unique variants given by parSEQ and NGS are justaposed in the two plots. **b)**The matching procedure showed that the majority of NGS reads correlated with high certainty to only one of the distinctive sequences. This was substantiated by noting that the Levenshtein distance ratios between the primary and secondary matches were predominantly beneath 0.5. **c)** Finally, the number of NGS reads for each unique parSEQ variant was plotted against the number of parSEQ hits (well support) for the distinctive read on a logarithmic scale. Note that NGS reads demonstrating a matching ratio exceeding 0.5 are not included in this plot.

The matching procedure showed that the majority of NGS reads correlated with high certainty to merely one of the distinctive sequences. This was substantiated by noting that the Levenshtein distance ratios between the primary and secondary matches were predominantly beneath 0.5, as shown in Figure 9,b. NGS reads demonstrating a matching ratio exceeding 0.5 were omitted. Subsequently, the number of NGS matches for each unique parSEQ read was plotted against the well support for the distinctive read on a logarithmic scale (Figure 9,c.

The derived plot elucidates that parSEQ sampling accurately mirrors the intrinsic pool distribution, thereby underscoring the precision and reliability of parSEQ in reflecting the genuine distribution of the original pool.

Eight out of the ten plates sampled with parSEQ ONT underwent new sequencing library preparations and were sequenced using the Illumina MiSeq Nano kit to facilitate a comparison of results (Illumina to ONT). We extracted 231 unique sequences from this run, aligning with our expectations. The sequences retrieved via Illumina sequencing matched those retrieved using ONT sequencing (results not shown).

Illumina sequencing results revealed superior accuracy when compared to ONT. This superiority was evident in the requirement of only tens of reads (approximately 50 Illumina reads per well) to establish an accurate well consensus sequence. ONT sequencing, on the other hand, necessitates hundreds of reads (at least 300 reads per well), to derive an accurate well consensus sequence.

Nevertheless, considering the substantially lower per-read-cost associated with ONT Minion in comparison to Illumina MiSeq, the adoption of ONT as the sequencing readout for parSEQ emerged as the more economical approach.

Unique sequences retrieved from this run were expressed in a PUREfrex 2.1 cell-free system ^26^ and screened for affinity against their antigen using bio-layer interferometry (BLI).

### 3.2 Case Study 2: parSEQ enables the retrieval of sequence verified, expression ready bacterial clones from bacterial pools

In the previous case study, we explored the use of parSEQ for the retrieval of unique small-binder-coding DNA variants from display pools. These retrieved variants were subsequently expressed in a PUREfrex 2.1 cell free system and screened. For such a workflow we used competent bacteria to transform plasmid pools holding the variants’ DNA. Post cherry-picking, the DNA template of every cherry picked variant was amplified from each well using high fidelity colony PCRs, and the non-purified DNA amplicant was subsequently used as template for PUREfrex expression of the protein variants. The pipeline was thus quite efficient as it permitted a fast turn around time from display pool to high numbers of expressed and sequence-verified protein variants ready for screening.

Using cell-free systems for the expression of such small binders (nanobodies, small synthetic binders) is appropriate as these binders have relatively simple structures. Moreover, we prefer working with cell-free systems when possible because they offer flexibility, speed, and cost-effectiveness when employed for protein expressions.

However, cell-free systems are not suitable for the expression of more complicated protein formats such as scFvs (single chain fragment variables) and Fabs (fragment antigen binding). While there has been studies reporting the successful expression of scFvs and Fabs in cell-free systems ^27,28,29,30,31^, our first hand experience indicates that cell-free systems are not reliable for the expression of these protein formats.

The second most simple system for expressing scFvs and Fabs is bacterial systems. ScFvs and Fabs have disulfide bonds that are crucial for the folding and stability of these proteins. In bacteria, disulfide bond formation predominantly occurs in the periplasmic space of gram-negative bacteria or the extracellular space in gram-positive bacteria, not in the cytoplasm ^32,33,34^. Thus, the most frequently used approach to produce scFvs and Fabs in E. coli is to express them in the periplasm of the bacteria. A newer approach though, employs E coli. SHuffle strains for the recombinant expression of proteins requiring the formation of disulfide bonds in the cytoplasm. E. coli shuffle strains have been engineered to have an oxidizing cytoplasm and to constitutively express a chromosomal copy of the di-sulfide bond isomerase DsbC, promoting the formation of di-sulfide bonds in the cytoplasm ^35,36,37^.

We chose to work with E. coli SHuffle B strain because it yielded better results concerning protein expression reliability. We transformed SHuffle B expression strain bacteria with a pET21a plasmid pool holding the scFv variant DNA. After transformation, the bacteria were plated in a lawn format and left overnight to grow. The following day, the lawn was scraped, re-suspended into growth media, and then fed into the parSEQ process.

Using parSEQ on a sampling of 4x 384-well plates, we were able to retrieve 268 unique variants from the scFv pool. These were cherry picked, transferred into a new 384 well plate, and stored at -80°C. Note that these are sequence verified, expression-ready, bacterial strains.

Chosen variants were then expressed in wells of deep 96-well plates. Expression was performed under a dual-temperature protocol after growing the bacteria in ThermoFisher’s auto-inducing MagicMedia. After expression, we implemented a thermal and osmotic shock protocol from Lindner et al. ^38^ for the fast purification of the expressed proteins in deep 96-well plates.

These sequence-verified, expressed, and purified (only partially purified under thermal and osmotic shock purification) scFvs were thus ready for affinity screening using BLI. The entire workflow is depicted in Figure 10.

**Figure 10:**
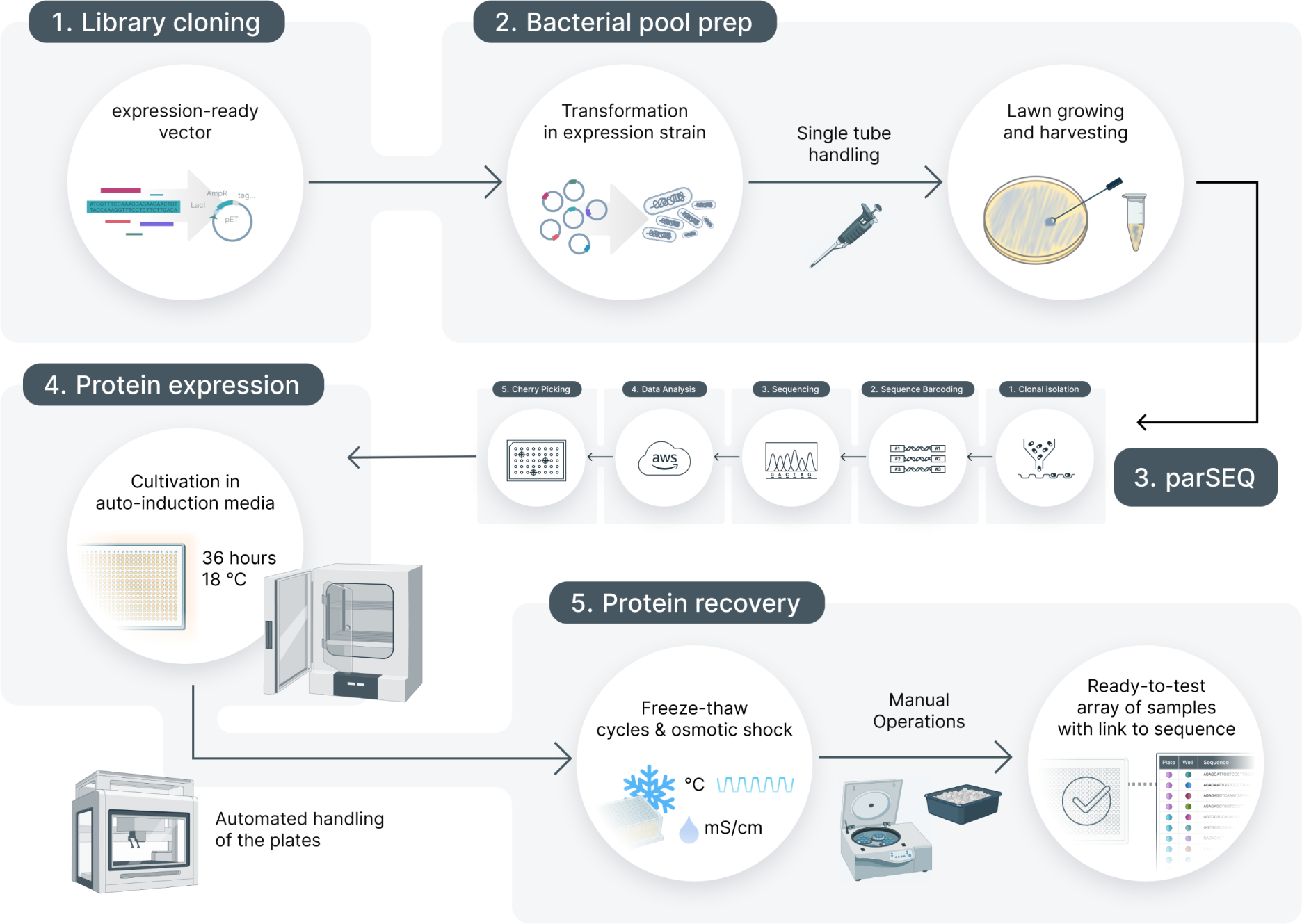
Case Study 2: parSEQ enables the fast isolation of expression-ready bacterial variants. An scFv library was first subcloned into an expression ready plasmid vector, then transformed into E.coli SHuffle expression strain. The transformation product was then plated in a lawn format on agar plates. The next morning, the lawn was scraped and resuspended into LB media, and the resulting pool was processed with parSEQ. Using parSEQ on a sampling of 4x 384-well plates, we were able to retrieve 268 unique sequences. Chosen variants were then expressed in wells of deep 96-well plates using auto-inducing MagicMedia under dual-temperature protocol. After expression, we implemented a thermal and osmotic shock protocol from Lindner et al. ^38^ for the fast purification of the expressed proteins in deep 96-well plates. These sequence-verified, expressed, and purified (only partially purified under thermal and osmotic shock purification) scFvs were thus ready for affinity screening using BLI.

### 3.3 Case Study 3: parSEQ enables lower-cost sourcing of DNA for testing denovo designed protein binders

Generative AI has proven to be a powerful tool for binder design, as demonstrated by the recent success of RFDiffusion in generating pico-molar binders without any experimental optimization ^39^. However, the thousands of potential binders generated by RFDiffusion must be experimentally evaluated to find the best-performing designs. This requires a tight feedback loop between the design and validation stages, ensuring a full Design-Test-Learn (DTL) cycle.

Historically, DTL cycles have been anchored in directed evolution strategies that are goal-driven, emphasizing the iterative enrichment of sequences to explore local maxima in the sequence-function landscape. Conversely, denovo generation techniques like RFDiffusion go beyond the traditional DTL cycle, and offer novel selection by exploring untapped and isolated regions of the landscape. An important challenge, however, is the procurement of DNA to experimentally validate these varied designs in a timely and cost-efficient manner.

We posited that parSEQ could help source DNA for denovo protein designs by integrating synthetic oligo libraries as a DNA source into the parSEQ process. Various DNA providers like Twist, Genscript, and IDT offer oligo libraries. IDT excels in providing premium quality, precisely normalized, and long oligo libraries (up to 350 bases), but at a higher price. Genscript, on the other hand, offers more affordable alternatives with shorter oligo lengths (up to 180 bases) and worse normalization ratios. We chose to work with Twist pools, accommodating long oligos (up to 300 bases) at competitive prices, albeit with suboptimal normalization ratios.

To showcase the ability of the pipeline, we designed a library of 1000 unique small binders employing denovo binder design strategies. We purchased a Twist oligo pool of 1000 synthetic oligos that code for therse binders. The oligo pool was then amplified and subsequently cloned into a plasmid backbone via Hifi cloning (Figure 11,a). Following transformation into competent bacteria, the library was sequenced using standard ONT MinIon sequencing. A sampling of 9x 384-well plates from the pool was also processed using parSEQ.

**Figure 11:**
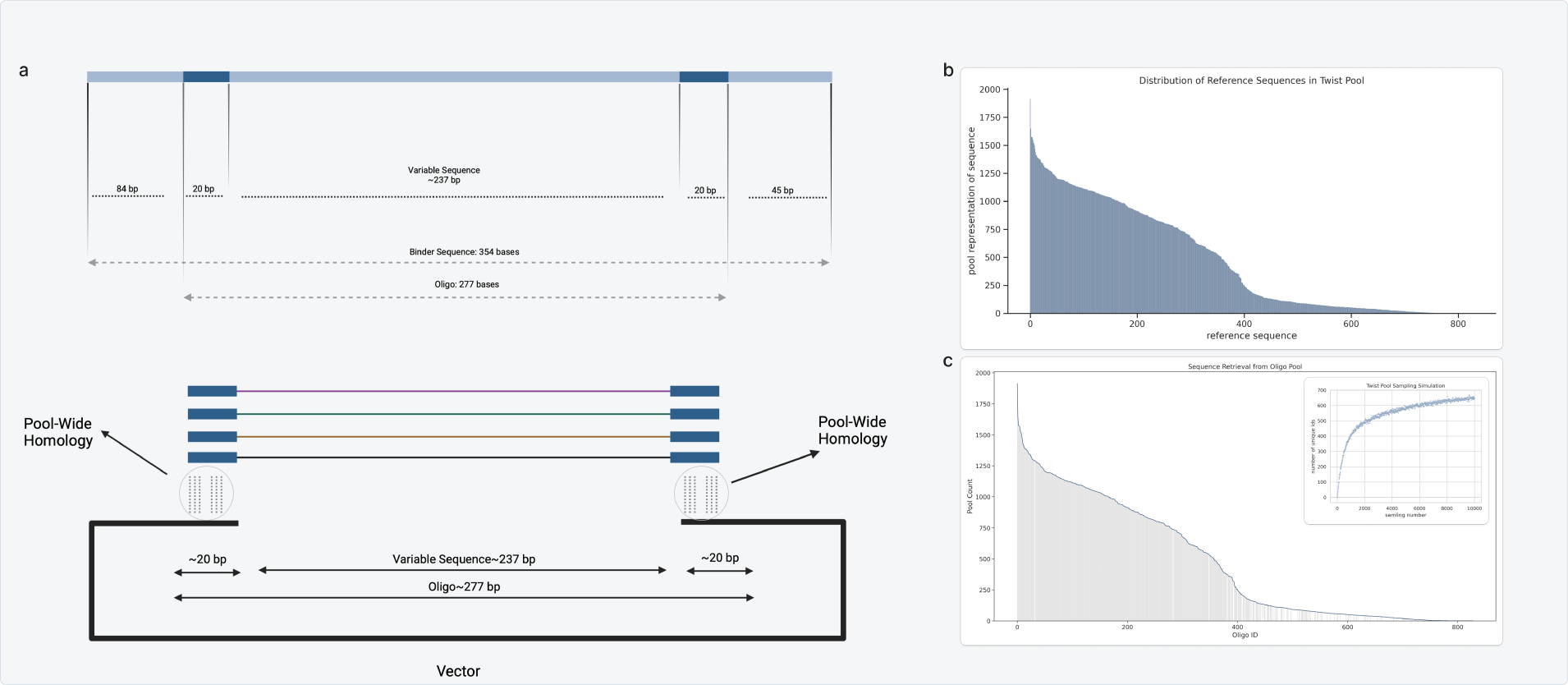
Case Study 3: parSEQ enables sourcing DNA for denovo-designed protein variants. A library of 1000 unique small binders were designed employing denovo binder design strategies. We purchased a Twist oligo pool of 1000 synthetic oligos that code for these binders. **a)** The oligo pool was amplified and subsequently cloned into a plasmid backbone via Hifi cloning. **b)** Following transformation into competent bacteria, the library was sequenced using standard ONT MinIon sequencing. Sequencing revealed the enrichment levels of the various designed binders in the pool. Evidently, substantial discrepancy exists in the enrichment of diverse designs within the Twist pool. 400 out of 1000 designs are well-represented, approximately 300 exhibit low representation, and the remaining 300 scarcely align with any of the 385,000 NGS reads. **c)** A sampling of 9x 384-well plates from the pool was also processed using parSEQ. We managed to recover 424 of the designed binders from the sample of 3,456 clones, aligning well with our sampling simulation based on the pool NGS distribution, when assuming a 0.85 parSEQ efficiency. The recovered oligos are superimposed over the pool enrichment profile. Predictably, variants exhibiting high pool enrichment were predominantly recovered.

NGS sequencing revealed the enrichment levels of the various designed binders in the pool (Figure 11,b). Evidently, a substantial discrepancy exists in the enrichment of diverse designs within the Twist pool—400 out of 1000 designs are well-represented, approximately 300 exhibit low representation, and the remaining 300 scarcely align with any of the 385,000 NGS reads.

It is important to highlight that microarrays yield pools of thousands of low concentration oligos and are usually prone to more errors than column-synthesized oligos ^40^. Moreover, microarray based manufacturing of oligo pools is notoriously susceptible to biases related to the synthesis efficiencies of varying oligos ^41^. Since these oligos are pooled directly without preceding normalization or quality control, the inherent synthesis biases pervade the pool. Additionally, the oligo pools supplied by Twist necessitate amplification before any substantial analysis can commence, due to the possibility of truncated oligos and the inherently low yield of microarray synthesis. This amplification process amplifies the already present bias in the pool and introduces new biases related to sequence amplification efficiency. These compounding biases and errors will lead to oligos being over- and under-represented in the pool.

Leveraging parSEQ, we managed to recover 424 of the designed binders from a sample of 3,456 clones, aligning well with our sampling simulation based on the pool NGS distribution, when assuming a 0.85 parSEQ efficiency. The recovered designs are depicted in Figure 11,c superimposed with the pool enrichment profile. Predictably, variants exhibiting high pool enrichment were predominantly recovered. This case study highlights the need for well-normalized, cost-effective pools. Access to such pools would significantly enhance our capacity to source DNA for proteins designed denovo.

### 3.4 Potential Application: parSEQ can potentially enable the retrieval of clonally verified sequences from massive site-directed mutagenesis libraries

Site-directed mutagenesis (SDM) studies are crucial in the fields of protein and genetic engineering. In SDM studies, researchers systematically introduce mutations at specific sites in a DNA sequence, enabling the study and manipulation of the sequence-function landscape of proteins. Site directed mutagenesis studies are important for understanding protein function, advancing enzyme engineering, drug development, protein-protein interaction studies, binding affinity maturation, functional genomics and proteomics studies, evolutionary studies, and as a general tool for synthetic biology ^42,43,44,45,46,47,48^.

For studies involving site-directed mutagenesis (SDM), scientists create libraries of variants by making precise changes to a DNA sequence. One popular method for preparing these variant libraries is called Polymerase Cycling Assembly (PCA) ^49,50,51^. PCA is a a variation of polymerase chain reaction (PCR) that utilizes DNA hybridization and annealing, as well as DNA polymerase, to assemble long DNA sequences from short oligos. Short DNA oligos are designed to hybridize together over overlapping regions. A DNA polymerase then extends and anneals the oligos to form the complete DNA strand. Finally, the completed DNA strand is amplified via PCR. (Figure 12, PCA Molecular Process).

**Figure 12:**
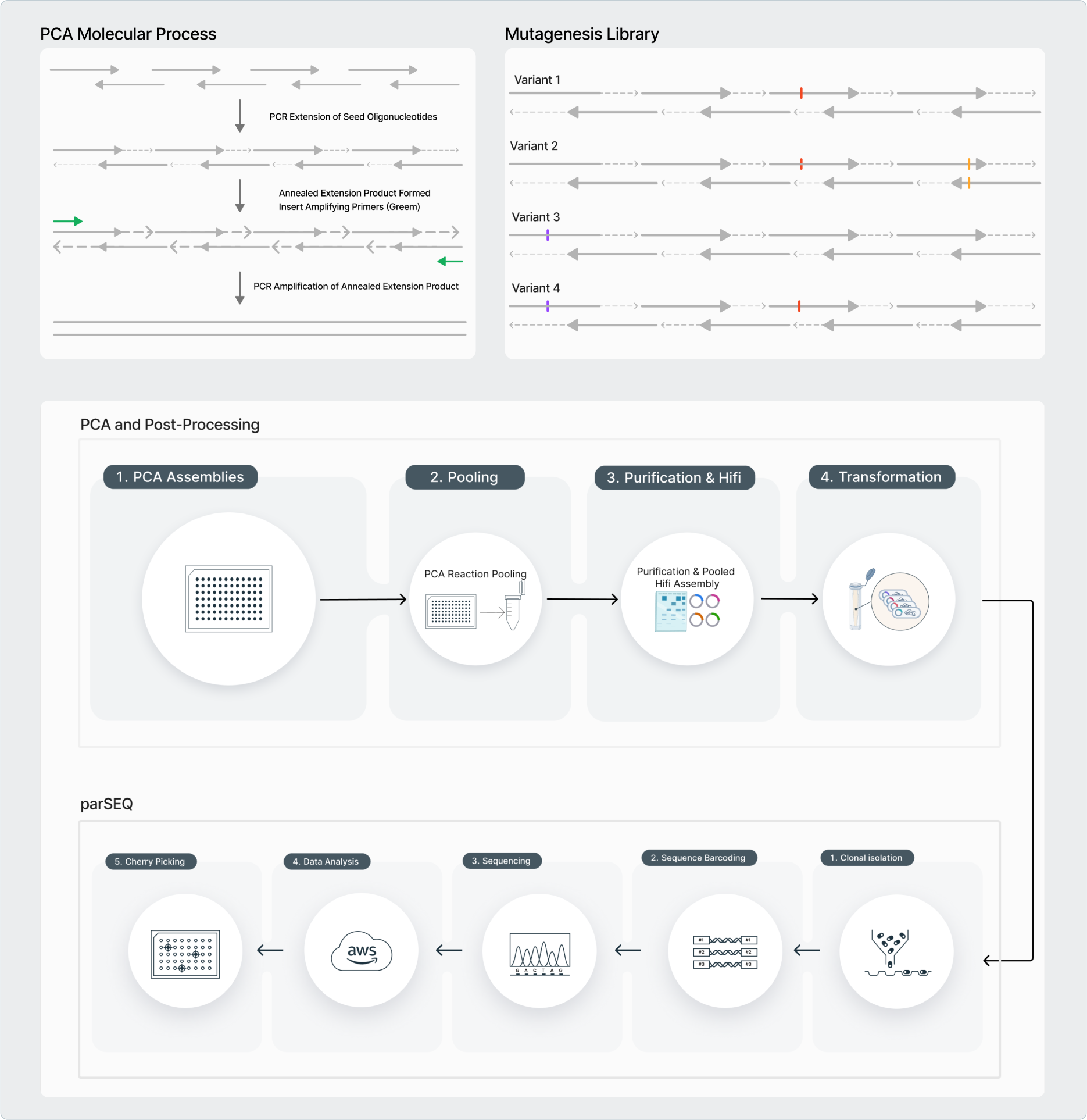
Case Study 4: parSEQ can potentially enable the retrieval of clonally verified sequences from massive site-directed mutagenesis libraries. **PCA Molecular Process:** Polymerase Cycling Assembly (PCA) is a a variation of polymerase chain reaction (PCR) that utilizes DNA hybridization and annealing, as well as DNA polymerase to assemble long DNA sequences from short oligos. **Mutagenesis Libraries:** Targeted mutagenesis libraries allow for a methodical examination of the effect of combinations of mutations on protein characteristics and activity. **PCA-parSEQ pipeline:** PCA assemblies are carried out for each variant separately in one well of a 384-well plate. The PCA reaction products are subsequently pooled and the pool is subcloned into a plasmid backbone via Hifi or Gibson Assembly. The circularized pool is then purified, transformed, and the transformed library is processed via parSEQ to retrieve sequence verified and physically separated variants of the mutational library.

PCA is advantageous because, when creating precise mutational libraries, the majority of oligonucleotides used during the design phase can be reused to generate the various variants, with the only requirement being the replacement of the oligos at the mutation sites (Figure 12, Mutagenesis Library). Consider a protein whose area of mutational study is 500 nts long. With oligos of 50 nts and an overlap of 20 nts, 14 oligos are required to assemble one variant. Therefore, to prepare 1200 possible variants, involving 60 possible mutation sites and their combinations, and assuming one mutation necessitates acquiring 1.5 new oligos (as some mutations will occur on oligo overlap regions), the original 14 oligos and an additional 90 oligos need to be purchased. The associated cost of acquiring these oligos is relatively low, slightly above $600, rendering Polymerase Cycling Assembly (PCA) a highly cost-effective approach for developing extensive parallel site-directed mutagenesis libraries.

Polymerase Cycling Assembly (PCA) comes with its own set of challenges though. Limitations include an inherent error rate due to the polymerase used, complexities causing misalignments and nonspecific bindings of oligos, and a resultant low yield, necessitating meticulous optimization. Often, researchers encounter these limitations in the form of low yield PCA reactions where a fraction of assembled DNA sequences contains errors ^52^.

In addition, the end product often needs to be in plasmid form, necessitating that the linear DNA product of PCA undergoes individual circularization for each reaction. Every plasmid, corresponding to a variant, needs subsequent bacterial transformation and sequence verification using Sanger sequencing.

We propose that parSEQ can resolve all all these limitations simultaneously.

After a standard PCA assembly where each variant is prepared in a separate well, all wells can be combined into one tube and purified collectively, before being circularized in a single, pooled Hifi Gibson-like assembly. This output can then be transformed into bacteria, and the resulting bacterial pool can be processed using parSEQ. After parSEQ processing, sequence-verified wells can be cherry-picked, allowing the retrieval of clonally verified variants (Figure 12, PCA and Post-Processing).

When comparing the overall PCA-parSEQ process to PCA followed by individual variant circularization, transformation, and Sanger-mediated sequence verification, the two approaches have similar timelines.

Concerning costs, the overall cost of retrieval of one sequence-verified variant (including the PCA assembly and labor cost) should be around $3-4 per variant. This is considerably cheaper than other options.

## 4 Conclusion

We have developed parSEQ to maximize the capture of sequence-function data-sets, ensuring that every screened variant’s functional data is paired with its sequence data. Operating within a 384-well plate setup, powered by Next-Generation Sequencing, and supported by automated procedures and Python-based data analysis, parSEQ can sample variant pools with high efficiency, low cost, and fast turnaround times. Moreover, parSEQ can be implemented in labs of varying scales, with low capital expenditure and a wide array of implementation options. parSEQ can be easily integrated into different workflows, as evident in our case studies. It can power the retrieval of physically separated and clonally verified variants from different DNA pool formats, including expression ready bacterial pools. It can also help in sourcing low-cost DNA for denovo designed proteins and for targeted mutational libraries. We recommend integrating parSEQ in the workflows of any team actively engaged in protein engineering, particularly those leveraging ML and/or AI.

## 5 Author Contributions

M.H. established the proof of concept, designed the pipeline, directed the development, and analyzed NGS data. F.P. performed most molecular biology experiments, developed molecular biology protocols, developed the parSEQ-PyHamilton Library, and analyzed NGS data. G.A. designed and built the infrastructure for analysis on the cloud and participated in setting up the automation. L.F. performed molecular biology experiments, and developed the expression-ready bacterial pool protocols. A.H. participated in establishing the Proof of Concept and performed molecular biology experiments. M.H. wrote the technical report with input from the remaining authors.

## 6 Code Availability

**parseq-analyze** is a python package that accompanies our parSEQ technical report. It is a python based pipeline to analyze parSEQ NGS results. This pipeline integrates multiple open-source packages and modules, with the main goal of identifying the consensus DNA sequence of every well that parSEQ investigates. parseq-analyze is available on the following GitHub repository: parseq-analyze.

**parseq-pyhamilton**, our automation library built on top of PyHamilton, is available on the following GitHub repository: parseq-pyhamilton. It contains our our core modules, as well as example protocols.

## References

1. Chenyi Li and Yajun Yan. Protein engineering for improving and diversifying natural product biosynthesis. In Trends in Biotechnology, doi: 10.1016/j.tibtech.2019.12.008, 2019.

2. Wenqian Li and Hafiz Iqbal. Broadening the Scope of Biocatalysis Engineering by Tailoring Enzyme Microenvironment: A Review. In Catalysis Letters, doi: 10.1007/s10562-022-04065-5, 2023.

3. Michaela Gebauer and Arne Skerra. Engineered Protein Scaffolds as Next-Generation Therapeutics. In Annual Review of Pharmacology and Toxicology, doi: 10.1146/annurev-pharmtox-010818-021118, 2020.

4. Kevin Yang and Frances Arnold. Machine-learning-guided directed evolution for protein engineering. In Nature Methods, doi: 10.1038/s41592-019-0496-6, 2019.

5. Harini Narayanan and Paolo Arosio. Machine Learning for Biologics: Opportunities for Protein Engineering, Developability, and Formulation. In Trends in Pharmacological Sciences, doi: 10.1016/j.tips.2020.12.004, 2021.

6. Bruce J. Wittmann and Frances H Arnold. Advances in machine learning for directed evolution. In Current Opinion in Structural Biology, doi: 10.1016/j.sbi.2021.01.008, 2021.

7. Bruce J. Wittmann and Frances H. Arnold. Informed training set design enables efficient machine learning-assisted directed protein evolution. In Cell Systems, doi: 10.1016/j.cels.2021.07.008, 2021.

8. Matthew S Faber and Timothy A Whitehead. Data-driven engineering of protein therapeutics. In Current Opinion in Biotechnology, doi: 10.1016/j.copbio.2019.01.015, 2019.

9. Adam McConnell and Benjamin J. Hackel. Protein engineering via sequence-performance mapping. In Cell Systems, doi: 10.1016/j.cels.2023.06.009, 2023.

10. Chase R Freschlin and Philip A Romero. Machine learning to navigate fitness landscapes for protein engineering. In Current Opinion in Biotechnology, doi: 10.1016/j.copbio.2022.102713, 2022.

11. Han Xiao, Zehua Bao, and Huimin Zhao. High Throughput Screening and Selection Methods for Directed Enzyme Evolution. In American Chemical Society, doi: 10.1021/ie503060a, 2014.

12. Norbert Furtmanna, Marion Schneider, and Joerg Birkenfeld. An end-to-end automated platform process for high-throughput engineering of next-generation multi-specific antibody therapeutics. In MABS, doi: 10.1080/19420862.2021.1955433, 2021.

13. Miten Jain, Nicholas J Loman, and Mathew Loose. Nanopore Sequencing and Assembly of a Human Genome with Ultra-Long Reads. In Nature Biotechnology, doi: 10.1038/nbt.4060, 2018.

14. Bruce J. Wittmann and Frances Arnold. evSeq: Cost-Effective Amplicon Sequencing of Every Variant in a Protein Library. In ACS Synthetic Biology, doi: 10.1021/acssynbio.1c00592, 2022.

15. Nicole Lerminiaux, Michael R. Mulvey, and Laura Mataseje. Do we still need Illumina sequencing data?: Evaluating Oxford Nanopore Technologies R10.4.1 flow cells and v14 library prep kits for Gram negative bacteria whole genome assemblies. In biorxiv, doi: 10.1101/2023.09.25.559359, 2023.

16. Thomas Sauvage, Alexandre Cormier, and Passerini Delphine. A comparison of Oxford nanopore library strategies for bacterial genomics. In BMC Genomics, doi: 10.1186/s12864-023-09729-z, 2023.

17. Shifu Chen and Jia Gu. fastp: an ultra-fast all-in-one FASTQ preprocessor. In Bioinformatics, doi: 10.1093/bioinformatics/bty560, 2018.

18. John A Hawkins and William H Press. Indel-correcting DNA barcodes for high-throughput sequencing. In PNAS, doi: 10.1073/pnas.1802640115, 2018.

19. Martin Šošić and Mile Šikić. Edlib: a C/C++ library for fast, exact sequence alignment using edit distance. In Bioinformatics, doi: 10.1093/bioinformatics/btw753, 2018.

20. Peter J. A. Cock and Michiel J. L. de Hoon. Biopython: freely available Python tools for computational molecular biology and bioinformatics. In Bioinformatics, doi: 10.1093/bioinformatics/btp163, 2009.

21. Robert C. Edgar. MUSCLE: multiple sequence alignment with high accuracy and high throughput. In Nucleic Acids Research, doi: 10.1093/nar/gkh340, 2004.

22. Kazutaka Katoh and Daron M. Standley. MAFFT Multiple Sequence Alignment Software Version 7: Improvements in Performance and Usability. In Molecular Biology and Evolution, doi: 10.1093/molbev/mst010, 2013.

23. Emma J Chory and Kevin M Esvelt. Enabling high-throughput biology with flexible open-source automation. In Molecular Systems Biology, doi: 10.15252/msb.20209942, 2021.

24. Weizhong Li and Adam Godzik. Cd-hit: a fast program for clustering and comparing large sets of protein or nucleotide sequences. In Bioinformatics, doi: 10.1093/bioinformatics/btl158, 2006.

25. Limin Fu, Beifang Niu, Zhengwei Zhu, Sitao Wu, and Weizhong Li. CD-HIT: accelerated for clustering the next-generation sequencing data. In Bioinformatics, doi: 10.1093/bioinformatics/bts565, 2012.

26. Jared L. Dopp and Nigel F. Reuel. Simple, Functional, Inexpensive Cell Extract for in vitro Prototyping of Proteins with Disulfide Bonds. In biorchiv, doi: 10.1101/2019.12.19.883413, 2020.

27. Satoshi Murakami, Rena Matsumoto, and Takashi Kanamori. Constructive approach for synthesis of a functional IgG using a reconstituted cell-free protein synthesis system. In Nature Scientific Reports, doi: 10.1038/s41598-018-36691-8, 2019.

28. Hanako Ishimaru and Yasuko Mori. Identification and Analysis of Monoclonal Antibodies with Neutralizing Activity against Diverse SARS-CoV-2 Variants. In Journal of Virology, doi: 10.1128/jvi.00286-23, 2023.

29. Takayasu Kawasaki and Yaeta Endo. Efficient synthesis of a disulfide-containing protein through a batch cell-free system from wheat germ. In European Journal of Biochemistry, doi: 10.1046/j.1432-1033.2003.03880.x, 2003.

30. Helmut Merk, Michael Gerrits, and Wolfgang Stiege. Cell-free synthesis of functional and endotoxin-free antibody Fab fragments by translocation into microsomes. In BioTechniques, doi: 10.2144/0000113904, 2018.

31. Helmut Merk and Volker A. Erdmann. Cell-Free Expression of Two Single-Chain Monoclonal Antibodies against Lysozyme: Effect of Domain Arrangement on the Expression. In The Journal of Biochemistry, doi: 10.1093/oxfordjournals.jbchem.a022290, 1999.

32. Katleen Denoncin and Jean-François Collet. Disulfide Bond Formation in the Bacterial Periplasm: Major Achievements and Challenges Ahead. In ARS Discoveries, doi: 10.1089/ars.2012.4864, 2013.

33. Cristina Landeta, Dana Boyd, and Jon Beckwith. Disulfide bond formation in prokaryotes. In Nature Microbiology, doi: 10.1038/s41564-017-0106-2, 2018.

34. Bruno Manta, Dana Boyd, and Mehmet Berkmen. Disulfide Bond Formation in the Periplasm of Escherichia coli. In EcoSal Plus, doi: 10.1128/ecosalplus.esp-0012-2018, 2019.

35. Julie Lobstein and Mehmet Berkmen. SHuffle, a novel Escherichia coli protein expression strain capable of correctly folding disulfide bonded proteins in its cytoplasm. In Microbial Cell Factories, doi: 10.1186/1475-2859-11-56, 2012.

36. Guoping Ren, Na Ke, and Mehmet Berkmen. Use of the SHuffle Strains in Production of Proteins. In Current Protocols in Protein Science, doi: 10.1002/cpps.11, 2016.

37. James B. Eaglesham, Augusto Garcia, and Mehmet Berkmen. Production of antibodies in SHuffle Escherichia coli strains. In Methods in Enzymology pages 105-144, doi: 10.1016/bs.mie.2021.06.040, Elsevier, 2021.

38. Robert Lindner and Martin Siemann-Herzberg. Process development of periplasmatically produced single chain fragment variable against epidermal growth factor receptor in Escherichia coli. In Journal of Biotechnology, doi: 10.1016/j.jbiotec.2014.10.003, 2014.

39. Joseph L. Watson and David Baker. De novo design of protein structure and function with RFdiffusion. In nature, doi: 10.1038/s41586-023-06415-8, 2023.

40. Ishtiaq Saaem and Jingdong Tian. In situ synthesis of DNA microarray on functionalized cyclic olefin copolymer substrate. In ACS Applied Material Interfaces, doi: 10.1021/am900884b, 2010.

41. Jason C. Klein, David Baker, and Jay Shendure. Multiplex pairwise assembly of array-derived DNA oligonucleotides. In Nucleic Acids Research, doi: 10.1093/nar/gkv1177, 2016.

42. Patricia E Carrigan, Petek Ballar, and Sukru Tuzmen. Site-directed mutagenesis. In Methods in Molecular Biology pages 107-124, doi: 10.1007/978-1-61737-954-3_8, Springer 2010.

43. Roberto A Chica, Nicolas Doucet, and Joelle N Pelletier. Semi-rational approaches to engineering enzyme activity: combining the benefits of directed evolution and rational design. In Current Opinion in Biotechnology, doi: 10.1016/j.copbio.2005.06.004, 2005.

44. Fei Wen, Michael McLachlan, and Huimin Zhao. Directed Evolution: Novel and Improved Enzymes. In Wiley Encyclopedia Of Chemical Biology, doi: 10.1002/9780470048672.wecb125, 2008.

45. Guangrong Xie and Jianhua Chen. Development of Therapeutic Chimeric Uricase by Exon Replacement/Restoration and Site-Directed Mutagenesis. In MDPI, doi: 10.3390/ijms17050764, 2016.

46. Eline Sijbesma and Christian Ottmann. Site-Directed Fragment-Based Screening for the Discovery of Protein–Protein Interaction Stabilizers. In Journal of the American Chemical Society, doi: 10.1021/jacs.8b11658, 2019.

47. Kiyoshi Yasukawa and Kuniyo Inouye. Improving the activity and stability of thermolysin by site-directed mutagenesis. In Biochimica et Biophysica Acta (BBA), doi: 10.1016/j.bbapap.2007.08.002, 2017.

48. Rafael Sanjuán. Mutational fitness effects in RNA and single-stranded DNA viruses: common patterns revealed by site-directed mutagenesis studies. In Phil. Trans. R. Soc., doi: 10.1098/rstb.2010.0063, 2010.

49. Carlos G. Acevedo-Rocha and Manfred T. Reetz. Assembly of Designed Oligonucleotides: A Useful Tool in Synthetic Biology for Creating High-Quality Combinatorial DNA Libraries. In Methods in Molecular Biology vol 1179 pages 189–206, doi: 10.1007/978-1-4939-1053-313, Springer 2014.

50. Ai-Sheng Xiong and Quan-Hong Yao. Chemical gene synthesis: strategies, softwares, error corrections, and applications. In FEMS Microbiology Reviews, doi: 10.1111/j.1574-6976.2008.00109.x, 2008.

51. Randall A. Hughes, Aleksandr E. Miklos, and Andrew D. Ellington. Chapter twelve - Gene Synthesis: Methods and Applications. In Methods in Enzymology, doi: 10.1016/B978-0-12-385120-8.00012-7, 2011.

52. Ai-Sheng Xiong and Quan-Hong Yao. Non-polymerase-cycling-assembly-based chemical gene synthesis: Strategies, methods, and progress. In Biotechnology Advances, doi: 10.1016/j.biotechadv.2007.10.001, 2008.

